# Mechanism of transcription modulation by the transcription-repair coupling factor

**DOI:** 10.1101/2021.04.02.438179

**Authors:** Bishnu P Paudel, Zhi-Qiang Xu, Slobodan Jergic, Aaron J Oakley, Nischal Sharma, Simon HJ Brown, James C Bouwer, Peter J Lewis, Nicholas E Dixon, Antoine M van Oijen, Harshad Ghodke

**Author notes:** Harshad Ghodke, Molecular Horizons and School of Chemistry and Molecular Bioscience, University of Wollongong, Wollongong NSW 2522, Australia.

## Abstract

Elongation by RNA polymerase is dynamically modulated by accessory factors. The transcription-repair coupling factor (TRCF) recognizes distressed RNAPs and either rescues transcription or initiates transcription termination. Precisely how TRCFs choose to execute either outcome remains unclear. With *Escherichia coli* as a model, we used single-molecule assays to study dynamic modulation of elongation by Mfd, the bacterial TRCF. We found that nucleotide-bound Mfd converts the elongation complex (EC) into a catalytically poised state, presenting the EC with an opportunity to restart transcription. After long-lived residence in this catalytically poised state, ATP hydrolysis by Mfd remodels the EC through an irreversible process leading to loss of the RNA transcript. Further, biophysical studies revealed that the motor domain of Mfd binds and partially melts DNA containing a template strand overhang. The results explain pathway choice determining the fate of the EC and provide a molecular mechanism for transcription modulation by TRCF.

## INTRODUCTION

Gene expression involves synthesis of RNA transcripts by RNA polymerases across all domains of life. During elongation, RNA polymerase (RNAP) maintains a 9 – 10 nucleotide RNA:DNA hybrid within the transcription bubble through the templated addition of ribonucleotides (reviewed by (1)). Processive nucleotide addition cycles result in the growth of the RNA transcript that is then extruded from the polymerase through the exit channel, accompanied by translocation of the polymerase along the template. Finally, during termination, nucleotide addition ceases and RNAP is displaced from the template DNA and recycled for subsequent rounds of transcription.

Elongation is punctuated by sequence-dependent pausing, stalling at sites of damage on the DNA template or at sites of roadblocks in the form of DNA-bound proteins. Prolonged residence in paused states can lead to escape into backtracked states in which the catalytic site of RNAP loses register with the 3’ end of the RNA transcript, allowing the newly synthesized transcript to invade the secondary channel of RNAP (2–4). Rescue from such backtracked states is catalyzed by cleavage factors such as GreA and GreB (2, 5). Elongation is modulated by the universally conserved elongation/pause factor, the bacterial NusG, homologous with Spt5 in eukaryotes (reviewed by (6)). In *Escherichia coli*, NusG maintains processive transcription (7) by suppressing backtracking through contacts on the upstream DNA (8, 9).

Elongation is also modulated by the transcription-repair coupling factor (TRCF) resulting in either transcription termination or rescue. The bacterial TRCF – Mfd is a multi-domain adaptor protein consisting of 7 domains connected by flexible linkers (Supplemental Figure 1A) (10). Domains 1a/2/1b form the UvrB − homology module that bears structural homology to the nucleotide excision repair protein UvrB and enables contacts with the repair protein UvrA. Domain 3 serves as a structural element playing a role in orienting the UvrB-homology module during the activation of the protein (11), following interactions of domain 4, the RNAP interaction domain, with the EC (12, 13). Domains 5 and 6 together form the motor domain, representing a (helicase) superfamily II ATPase that exhibits structural homology to the RecG translocase (10, 14). Domain 7 is an auto-inhibitory domain that contacts the UvrB-homology module to maintain it in a closed conformation, suppressing promiscuous DNA binding of Mfd in the absence of interactions with RNAP.

A detailed understanding of the activities of Mfd has been assembled through a combination of molecular biology, biochemical (13,15–21), structural methods (10-12,22), as well as single-molecule (23–26) and live-cell imaging techniques (27–29). Mfd acts in a complex role in modulating transcription activity, including at hard-to-transcribe regions (30). At sites of DNA damage, Mfd orchestrates premature transcription termination leading to an enhancement in repair of DNA damage in the template strand − a phenomenon known as transcription-coupled DNA repair (16,21,26,28,29,31). In bacterial cells, the UvrA_2_B nucleotide excision repair machinery is recruited to the site of distressed RNAPs by Mfd, resulting in the site-specific loading of the damage verification enzyme UvrB (28, 29). This ability to recruit DNA-repair machinery at sites of lesion-stalled RNAP to promote ATP-dependent premature transcription termination is a property that is shared by eukaryotic functional homologs of Mfd – the yeast Rad26 (32–34) and human CSB (35, 36). Mutations in CSB protein result in deficient repair machinery recruitment causing Cockayne syndrome in humans (36–38).

In addition to initiating transcription termination, the three best-studied TRCFs also serve as elongation factors. Like Rad26 (32) and CSB (39), Mfd has also been shown to rescue transcription at pause sites *in vitro* through a ‘release-and-catch up’ mechanism (13, 24). Forward translocation of the EC is central to TRCF activities including rescue from backtracked states and transcription termination (13,24,32,39). Considering their membership in the class of superfamily 2 helicases (40), the structural homology of the TRCF motor domains and a similar mode of engagement with the EC between Mfd (11,12,22) and Rad26 (32), it is plausible that the molecular mechanisms underlying these outcomes are conserved.

Precisely how interactions of TRCF with the cognate EC result in transcription termination versus rescue is unknown. These interactions can be thought of as proceeding via three stages. First, during initial recognition, the TRCF binds an EC through a direct protein-protein interaction (32). Second, the TRCF is loaded on to the EC in a step that may require ATP hydrolysis and translocation on the upstream DNA up to the EC (12,22,32). Finally, in the third, remodeling stage, correctly loaded TRCF remodels the EC to modulate an outcome either by itself or by recruiting additional factors to promoting transcription elongation or termination (23,24,32,41).

Two models have been proposed to explain forward translocation and remodeling of RNAP by TRCFs: first, the TRCF ‘pushes’ RNAP forward while translocating on dsDNA as proposed for Mfd (12, 13), and second, the TRCF pulls the ssDNA template to effectively translocate the polymerase forward (32). Recent structures of Mfd loading on an *in vitro* reconstituted EC assembled on a non-complementary bubble have demonstrated that following initial recognition of the EC, Mfd translocates forward until it bumps into the RNAP (12). Considering that Mfd translocates by one nucleotide per round of ATP hydrolysis, a hypothetical model for the mechanism of Mfd-mediated collapse of the transcription bubble has been proposed in which Mfd rewinds the transcription bubble by pulling double-stranded DNA one base pair at a time through forward translocation (12). In this model (Model I), it is proposed that torque exerted by Mfd leads to positive supercoiling on the DNA enabling reannealing of the transcription bubble and displacement of the transcript, and along with it, RNAP. Direct supporting evidence in the form of step-wise rewinding of the transcription bubble is not currently available.

Eukaryotic homologs of Mfd are proposed to employ a different mechanism (Model II). The structure of a remodeling intermediate formed by yeast Rad26 bound to upstream DNA in the RNA Pol II EC (32) reveals an extended transcription bubble and direct contacts between dsDNA-bound Rad26 and the ssDNA template strand. Based on the homology to the Swi2/Snf2-family core ATPase domains, and the mechanism of Snf2 in chromatin remodeling, it has been proposed that Rad26 pulls template DNA in the extended transcription bubble of Pol II EC (32). Experimental support for this mechanism of action for Mfd is insufficient, but it is notable that the motor domains of Mfd bear structural homology to motor domains of ATPases involved in chromatin remodeling (most notably Snf2) and its eukaryotic counterparts in transcription-coupled DNA repair (11).

Since available structures depict initial loading of Mfd on the EC but not transcription termination leading to disassembly of the EC accompanied by loss of the transcript (12), and given that Mfd shares significant structural homology with chromatin remodeling factors (11), the molecular mechanism underlying rewinding of the transcription bubble and transcription termination is not yet a settled question. Further, while it is clear that displacement of the polymerase in the absence of downstream binding partners of Mfd is a slow event requiring several minutes *in vitro* (23, 25), the sequence of events leading to loss of the transcript and its residence time in the EC during this remodeling remains unclear. We considered the two described models in detail and summarized predictions from theme: In the case of Model I, Mfd bound to the EC pulls dsDNA through, causing rewinding of the transcription bubble. A key prediction of this model is that an assay that is sensitive to single base-pair changes in the architecture of the transcription bubble with sufficiently high temporal resolution should be able to directly detect step-wise rewinding of transcription bubble, one nucleotide at a time. Since Mfd-mediated transcription termination requires several seconds *in vitro*, sub-second resolution should be sufficient to detect these step-wise conformational changes while rewinding the 9 – 10 nt transcription bubble. In contrast, a prediction of Model II is that binding of Mfd must lead to melting of upstream DNA, and Mfd must make contact with either the template or the non-template ssDNA. The residue(s) in contact must possess the ability to translate conformational changes occurring from cycles of ATP hydrolysis into a ‘pulling’ motion where it pulls ssDNA.

Here, we set out to understand how Mfd dynamically remodels the EC by monitoring the fate of the RNA transcript under various scenarios. We developed a single-molecule Förster resonance energy transfer (smFRET)-based assay that is sensitive to changes in the architecture of the transcription bubble at the level of single base-pairs by directly visualizing dynamic changes in the distance between donor-labeled upstream DNA and acceptor-labeled RNA transcript forming the transcription bubble. We reconstituted catalytically-functional ECs *in situ* and monitored the FRET signal from surface-immobilized molecules under conditions of partial or complete nucleotide starvation, and probed global remodeling of the EC by NusG and Mfd on the timescale of minutes. NusG efficiently rescued ECs from catalytically inactive states, consistent with its role as an elongation factor. Mfd was able to rescue ECs from catalytically inactive states into a long-lived catalytically poised state, prior to continued ATP-hydrolysis-dependent forward translocation leading to transcription termination. This catalytically poised state acts as a ‘molecular timer’, presenting the EC with an opportunity to restart transcription. Further biophysical and structural biology studies reveal that the nucleotide-bound motor domain melts the dsDNA on DNA substrates mimicking the transcription bubble and uncover a novel role for a conserved arginine residue as a “hook” in mediating contacts with template strand ssDNA. Collectively, the studies enable us to propose a ‘melt, hook and pull’ model to rationalize how ATP binding and hydrolysis by Mfd modulate transcription elongation or termination.

## MATERIAL AND METHODS

### Proteins used in this study

#### Purification of RNA polymerase

His-tagged RNA polymerase was purified essentially as described previously (42). pIA900 was a generous gift from Prof. Irina Artsimovitch (Addgene #104401)(42). pIA900 was transformed into BL21(λDE3)*recA*^-^ cells. A single colony was isolated and streaked on an ampicillin plate (100 mg L^-1^). Four clones were then tested for small scale protein expression (25 mL) and were found to produce similar levels of overexpressed RNAP. One colony was then used for overproduction in 6 L of LB medium, inoculated with an overnight culture and shaken for 2.5 h at 37 °C to reach *A*_595_ = 0.6, followed by induction with a final concentration of 1 mM IPTG for an additional 3 h, followed by pelleting. Overproduced protein was purified from cells following the exact protocols described in (42). Following chromatographic separation over the Mono-Q column, two fractions were isolated, one corresponding to the RNAP core complex, and a second smaller fraction corresponding to the holoenzyme. For this work, the RNAP core complex was isolated at a concentration of 35.7 mg mL^-1^ corresponding to 92 µM in storage buffer (50 mM Tris/HCl pH 7.6, 100 mM NaCl, 1 mM EDTA, 1 mM dithiothreitol and 50% *v*/*v* glycerol). Purified protein was stored at –80 °C following snap freezing with liquid N_2_. Aliquots in use were stored at –20 °C.

#### Overproduction and purification of Mfd variants

Sequence verified *mfd-ypet, mfd(E730Q)-ypet* and *mfd(R953A)-ypet* genes were obtained from Aldevron. Plasmids for overexpression of Mfd, Mfd(E730Q) and Mfd(R953A) were created as follows: first, *mfd* or mutant gene inserts (lacking the *ypet* gene) were amplified using PCR from the custom synthesized constructs using the primers (pETMCSII_Mfd_F and pETMCSII_Mfd_R; Supplemental Table 1). PCR products were digested with with *Bam*HI and *Eco*RI and purified using a Qiagen miniprep kit. The *mfd* (or mutant) fragments were inserted between the same restriction sites in vector pSH1017 (43), previously isolated using electrophoretic separation and an extraction from a 1% (*w*/*v*) agarose gel. Ligations were performed with T4 DNA ligase at 16 °C overnight, followed by transformation into *E. coli* DH5α cells; colonies were screened for the presence of insert by colony PCR and intact *mfd* sequences in plasmids pHG4137 (coding for wildtype Mfd), pHG42305 (Mfd(R953A)) and pHG42306 (Mfd(E730Q)) were confirmed by sequencing.

A generalized procedure for the expression and purification of Mfd variants used in this study is best exemplified by the methods used in the production of Mfd(E730Q). *E. coli* strain BL21(λDE3)*recA*/pHG42306 (E730Q) was grown at 37°C in LB medium supplemented with thymine (50 mg L^-1^) and ampicillin (200 mg L^-1^) to *A*_600_ = 0.6. To induce Mfd(E730Q) overproduction, 1 mM IPTG was added to the shaking culture. Cultures were grown for a further 3 h, and then chilled in ice. Cells were harvested by centrifugation (11,000 x *g*; 6 min), frozen in liquid N_2_ and stored at –80 °C.

After thawing, cells (17.1 g, from 6 L of culture) were resuspended in 120 mL lysis buffer (50 mM Tris/HCl, pH 7.6, 2 mM dithiothreitol, 0.5 mM EDTA, 20 mM spermidine) and lysed by two passages through a French press operated at 12000 psi. Cell debris was removed from the lysate by centrifugation (35,000 x *g*; 30 min) to yield the soluble Fraction I. Proteins in Fraction I that were then precipitated by addition of solid ammonium sulfate (0.38 g mL^-1^), followed by stirring for 30 min, were collected by centrifugation (38,000 x *g*; 30 min) and dissolved in 45 mL buffer A (50 mM Tris/HCl, pH 7.6, 2 mM dithiothreitol, 0.5 mM EDTA, 10% *v*/*v* glycerol) containing 180 mM NaCl. The solution was dialyzed against 2 L of the same buffer to yield Fraction II.

Fraction II was divided into two sub-fractions (23 mL each), which were successively applied at 1 mL min^-1^ onto a column (2.5 x 15 cm) of Toyopearl DEAE-650M resin that had been equilibrated in buffer A + 190 mM NaCl. Fractions containing proteins that did not bind to the resin (50 mL per sub-fraction; 100 mL in total) were pooled and dialyzed against two changes of 2 L of buffer A to yield Fraction III.

Fraction III was loaded at 1 mL min^-1^ onto the same DEAE column that had been equilibrated with buffer A. After the column was washed with 80 mL of the same buffer, Mfd(E730Q) was eluted at 2.5 mL min^−1^ using a linear gradient (700 mL) of 0−500 mM NaCl in buffer A. The Mfd(E730Q) eluted in a single, relatively narrow, peak centered around 80 mM NaCl. Fractions containing Mfd(E730Q) (40 mL) were pooled and dialyzed against four changes of 1.2 L of buffer B (50 mM Na/MES, pH 6.5, 2 mM dithiothreitol, 0.5 mM EDTA, 10% *v*/*v* glycerol) to yield Fraction IV.

Fraction IV (40 mL) was loaded at 1.5 mL min^-1^ onto a SP Sepaharose FastFlow column (Cytiva; 2.5 x 13 cm) that had been equilibrated with buffer B. After washing the column with 80 mL of buffer B, Mfd(E730Q) was eluted at 2.5 mL min^-1^ using a linear gradient (700 mL) of 0−500 mM NaCl in buffer B. The Mfd(E730Q) eluted in a single peak centered around 100 mM NaCl. Fractions containing highly purified Mfd(E730Q) were pooled and dialyzed against four changes of 1.2 L of storage buffer (40 mM Tris/HCl, pH 7.6, 100 mM NaCl, 4 mM dithiothreitol, 0.5 mM EDTA, 30% *v*/*v* glycerol) to yield final product containing 130 mg of protein at a final concentration of 46.7 µM. The same protocol yielded 51.5 mg of Mfd(R953A) at a final concentration of 18 µM. Final protein preparations were snap frozen in liquid N_2_ and stored at –80 °C.

In case of wildtype Mfd, a modified purification procedure was used whereby the fractions containing Mfd from the second DEAE (40 mL) column were pooled and directly applied at 1 mL min^-1^ onto a 2.5 x 10 cm column of heparin Sepharose 4B that had been equilibrated with buffer A + 50 mM NaCl. Mfd was eluted at 1 mL min^−1^, in a broad peak centered around ∼120 mM NaCl, using a linear gradient (450 mL) of 50−400 mM NaCl in buffer A. Fractions containing highly purified Mfd were pooled and dialyzed against 2 L of storage buffer (40 mM Tris/HCl, pH 7.6, 100 mM NaCl, 4 mM dithiothreitol, 0.5 mM EDTA, 25% *v/v* glycerol) to yield 115 mg of protein at a final concentration of 40 µM. Purified Mfd was snap frozen in liquid N_2_ and aliquots were stored at –80 °C. NusG was a generous gift from Prof Robert Landick (University of Wisconsin-Madison) (9).

#### Oligonucleotides and DNA substrates

All oligonucleotides were purchased from Integrated DNA Technologies (IDT, Singapore). Sequences are presented in Supplemental Table 1.

#### Single molecule FRET measurements

##### smFRET TIRF microscope setup

A home built objective-type TIRF microscope based on a Nikon TiE2 model was used to record single molecule movies. FRET was measured by excitation with a 532 nm laser and the emissions at 555 and 637 nm were collected using a band-pass filter at 555 nm and a long-pass filter at 650 nm. Scattered light was removed by using a 560 nm long pass filter. Cy3 and Cy5 signals were separated by a 638 nm dichroic using photometrics dual view (DV-2) and both signals were focused onto a CCD camera (Hamamatsu C9 100–13), simultaneously. Data were collected at 200 ms (5 frames per s) time resolution. A power density between 50−150 W cm^-2^ was used in typical experiments.

##### Preparation of flow cells for imaging

Coverslips were prepared as described previously (44). Quartz coverslips were cleaned with 100% ethanol and 1 mM KOH. Aminosilanization of cleaned coverslips was carried out using a 1% (*v*/*v*) (3-aminopropyl)triethoxy silane (Alfa Aesar, A10668, UK) solution in acetone. PEGylation was carried out by incubating a mixture of biotinPEG-SVA MW 5000 and mPEG-SVA MW 5000 (Laysan Bio, AL) at a ratio of 1:10 prepared in 50 mM MOPS pH 7.5 solution on the top of the silanized coverslip for 3–4 h. PEGylated coverslips were stored under dry nitrogen gas at −20 °C.

Neutravidin solution was prepared in transcription buffer (1 × TB; 20 mM Tris/HCl pH 8.0, 40 mM KCl, 5 mM MgCl_2_ and 1 mM β-mercaptoethanol) and spread on the top of dry a PEGylated coverslip followed by a 10 min incubation. Sample flow chambers were created by sandwiching polydimethylsiloxane (PDMS) on the top of the streptavidin coated coverslip. Then, blocking buffer (1 × TB + 0.25% (*v*/*v*) Tween 20) was injected into the channel to reduce non-specific binding of proteins on the surface followed by 10–15 min incubation. This was then followed by washing with 1 × TB to remove unbound Tween 20.

#### *In situ* reconstitution of ECs for smFRET measurements

Protocols described by (45) were adapted for *in situ* reconstitution of ECs in single-molecule FRET studies. Template DNA (1 µM) was annealed to RNA primer (3 µM) in 1 × TB by heating to 65 °C for 5 min followed by 3 °C drops in temperature every 2 min until the reaction temperature reached 27 °C, followed by a hold step at 25 °C. Typically, 200 µL of reaction were prepared at a time. To form the RNAP:template (T70_31Cy3):RNA(R15_4Cy5) complex, 0.22 µL of 92 µM RNAP was incubated with 2 µL of template:RNA hybrid in 1 × TB at room temperature for 30 min. Meanwhile, 200 µL of 100 pM of non-template NT70_bio (Supplemental Table 1) were immobilized on the neutravidin-coated surface. Following this, the RNAP:template:RNA mixture was diluted serially 6000 × in a final suspension of 200 µL of 1 × TB and introduced into the flow cell and incubated for at least 5 min to promote annealing of the template DNA to the biotinylated non-template DNA resulting in formation of the EC. At the end of this incubation step, the flow cell was washed with 200 µL of 1 × TB + 1 M KCl and subsequently with 1 × TB supplemented with an oxygen-scavenging system (OSS) consisting of protocatechuic acid (2.5 mM) and protocatechuate-3, 4-dioxigenase (50 nM) to reduce photo-bleaching of the fluorophores and 2 mM Trolox to reduce photo-blinking of dyes, followed by imaging. All elongation experiments in Figures 1 and 2 were performed at 100 µM of each NTP or dNTP (where indicated). smFRET experiments involving NusG were performed at 500 nM of NusG in 1 × TB (Figure 2). smFRET experiments involving wildtype Mfd (Figures 3 and 4) were performed at 500 nM of Mfd in 1 × TB. Experiments involving Mfd(R953A) (Figure 3 and 4) and Mfd(E730Q) (Figure 4) were performed at 1 µM of Mfd mutant in 1 × TB. Concentrations of ATP and ATPɣS were at 1 mM and 0.5 mM respectively unless otherwise specified.

**Figure 1:**
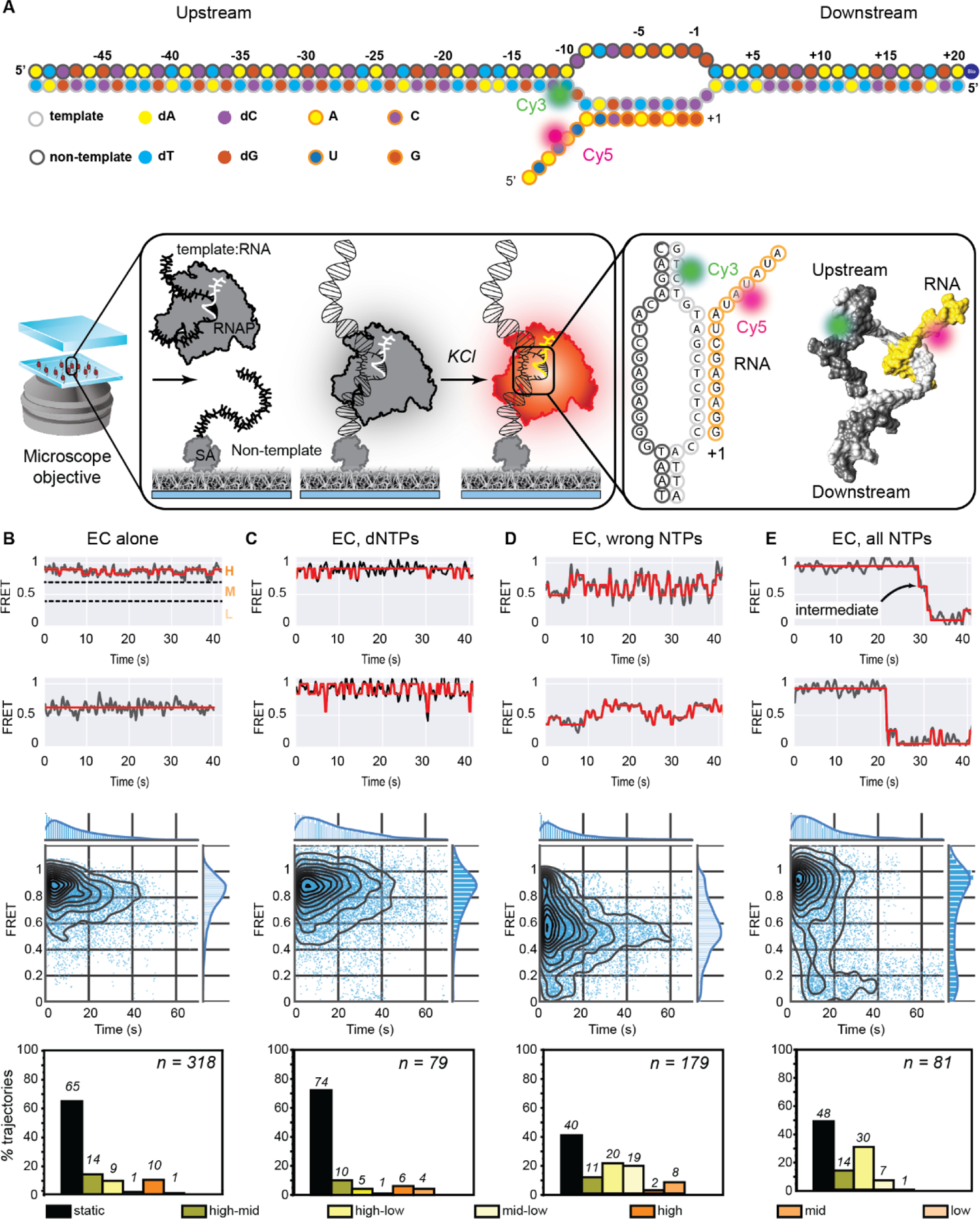
Reconstitution of a functional EC. (A) Schematic of the transcription bubble used in this study. The sequence of the four next correct incorporations is G, U, A and A. ECs were assembled on dye-labeled nucleic acids by flowing in pre-incubated RNAP-template:RNA complexes into a flow cell with immobilized biotinylated non-template strand (EC shown in grey). Following initial immobilization, a high-salt wash step was used to remove poorly hybridized ECs from the surface. Here, only complexes containing both the Cy3-labeled template strand (grey circles) and Cy5-labeled transcript (orange circles) are observed when stably associated with the biotinylated non-template strand (black circles). (B-E) Two example FRET trajectories (black) and idealized fits (red) (top panel), temporal heat maps (middle panel) and column plots indicating percentages of trajectories exhibiting static or dynamic behavior (exhibiting transitions between or within high, mid, or low FRET regimes) (bottom panel) are provided for (B) ECs alone (n = 318 molecules) or in the presence of (C) dNTPs (n = 79 molecules), (D) non-complementary (‘wrong’: UTP, ATP, CTP) NTPs (n = 179 molecules) and (E) the full set of NTPs (n = 81 molecules) respectively.

**Figure 2:**
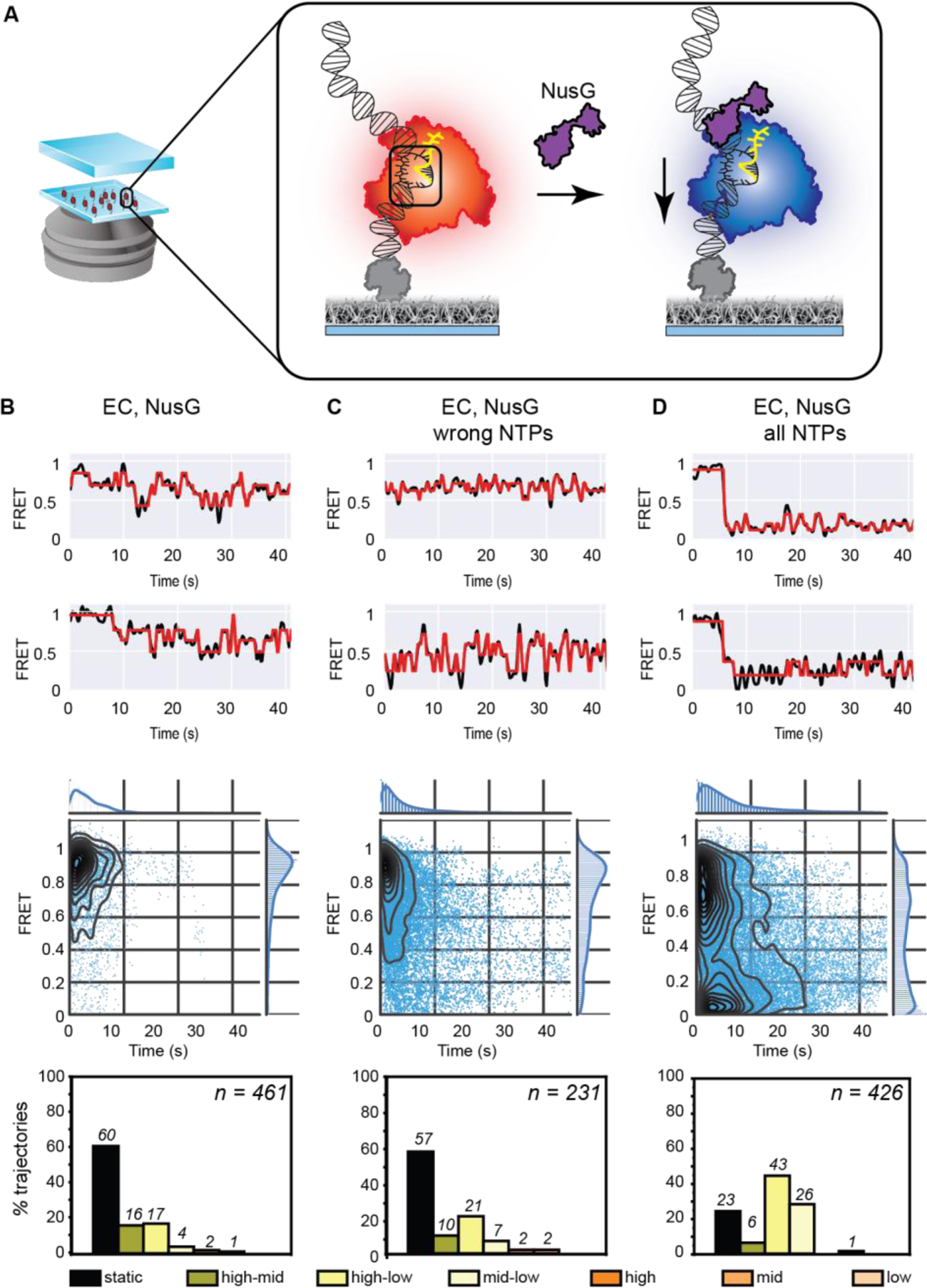
Interactions between NusG and the EC. (A) Schematic of FRET-pair labeled ECs incubated with NusG (purple). (B-D) Two example FRET trajectories (black) and idealized fits (red) (top panel), temporal heat maps (middle panel) and column plots indicating percentages of trajectories exhibiting static or dynamic behavior (exhibiting transitions between or within high, mid, or low FRET regimes) (bottom panel) are provided for (B) EC and NusG alone (n = 461 molecules), (C) EC and NusG in the presence of all wrong NTPs (n = 231 molecules), and (D) EC and NusG with the full set of NTPs (n = 426 molecules). NusG promotes the residence of the EC in a catalytically poised state, and enhances transitions to low FRET states.

**Figure 3:**
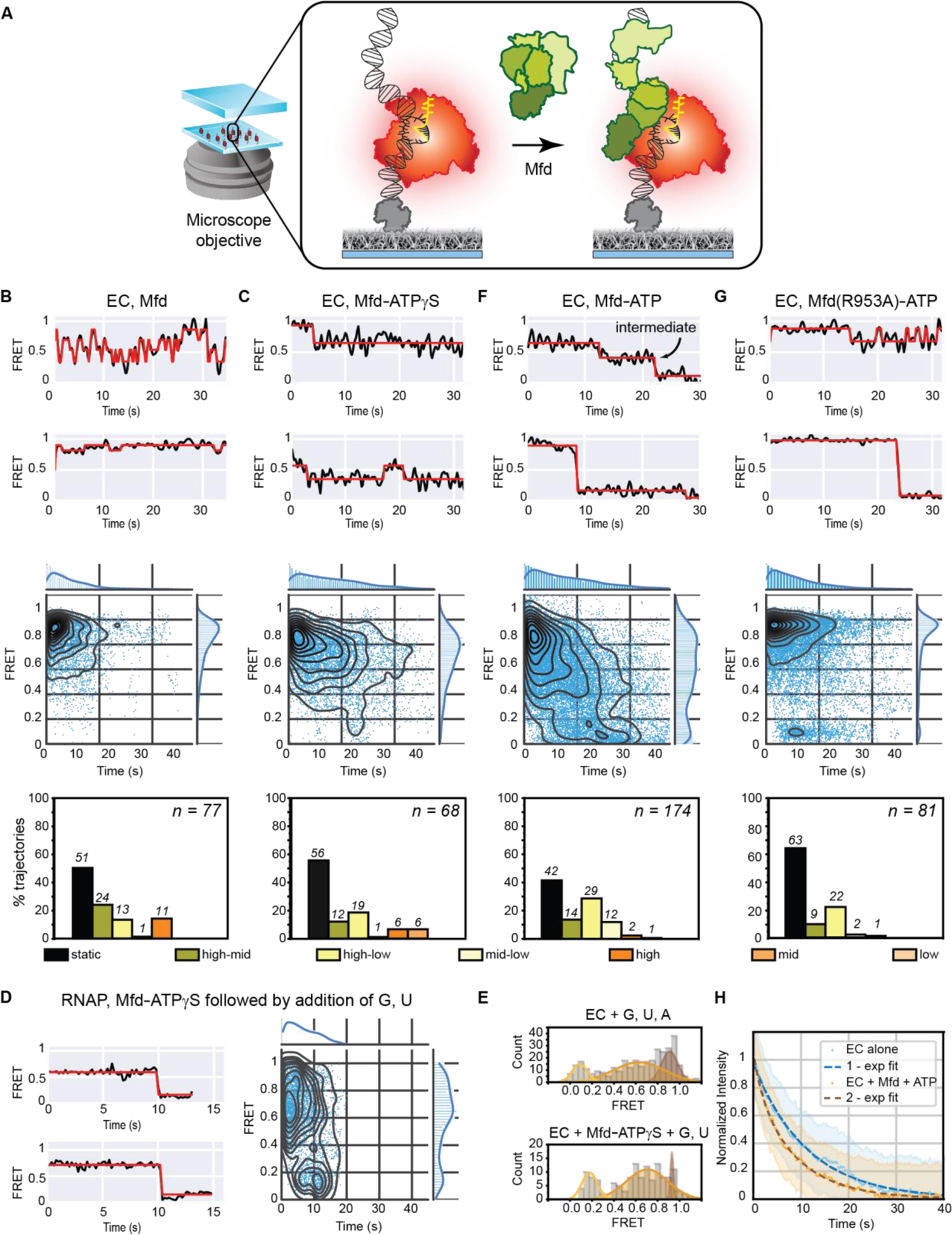
Interactions between Mfd and the EC. (A) Schematic of FRET-pair labeled, surface-immobilized ECs incubated with Mfd in solution. (B-C) Two example FRET trajectories (black) and idealized fits (red) (top panel), temporal heat maps (middle panel) and column plots indicating percentages of trajectories exhibiting static or dynamic behavior (exhibiting transitions between or within high, mid, or low FRET regimes) (bottom panel) are provided for (B) EC and Mfd alone (n = 77 molecules), (C) EC and Mfd in the presence of 0.5 mM ATPɣS (n = 68 molecules). (D) Example FRET trajectories of ECs incubated with Mfd and ATPɣS followed by addition of 100 μM each of GTP and UTP. Corresponding heat map shows an overview of the transition to low-FRET states upon incorporation of GTP and UTP. (E) Histograms (gray bars) of FRET states accessed by the EC when incubated with 100 μM each of GTP, UTP and ATP (top panel), or pre-incubated with Mfd-ATPɣS (10 min) followed by 100 μM each of GTP and UTP. Each histogram was fit to the sum of three individual Gaussian curves (see Supplemental Note 3). (F-G) Two example FRET trajectories (black) and idealized fits (red) (top panel), temporal heat maps (middle panel) and column plots indicating percentages of trajectories exhibiting static or dynamic behavior (exhibiting transitions between or within high, mid, or low FRET regimes) (bottom panel) are provided for (F) EC and Mfd in the presence of 1 mM ATP (n = 174 molecules) and (G) EC and DNA-binding competent but translocase-deficient Mfd(R953A) in the presence of 1 mM ATP (n = 81 molecules). (H) Normalized intensity of Cy5 fluorescence in the field of view following direct red laser excitation of the EC alone (blue circles) or incubated with Mfd and 1mM ATP (yellow circles). Single (blue) and double (brown) – exponential fits to the EC alone or when incubated with Mfd and 1 mM ATP are indicated as dashed lines. Error bars represent standard error of the fit.

**Figure 4:**
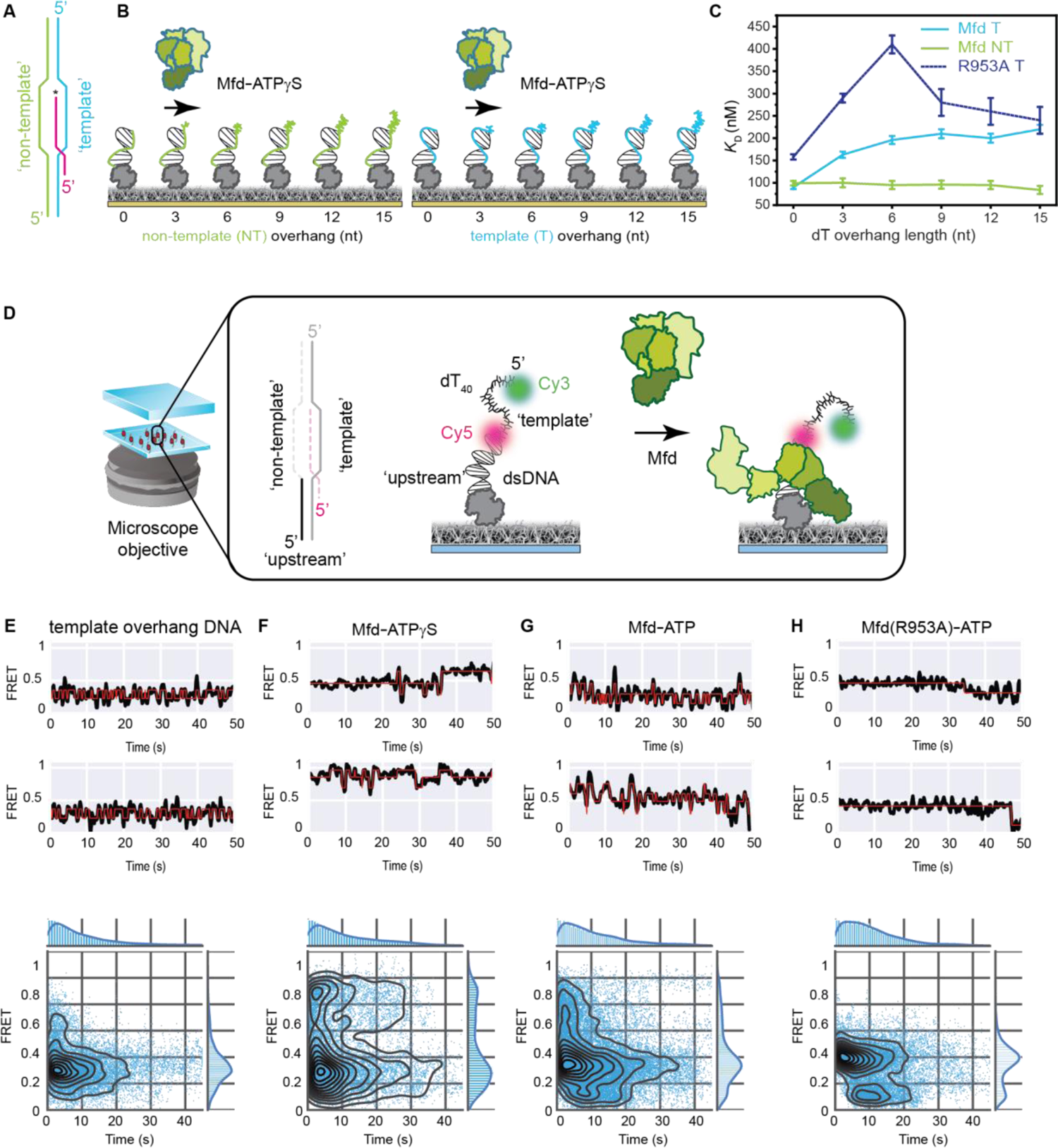
Interactions between Mfd and a primed-DNA template. (A) Schematic of a transcription bubble. (B) Schematic of SPR experiments to assess the strength of interaction between Mfd-ATPɣS or R953A-ATPɣS and primed DNA templates with 18 bp dsDNA and either non-template or template strand overhangs (dT_3_, dT_6_, dT_9_, dT_12_ or dT_15_ overhangs). (C) Plot depicting *K*_D_ of interaction as a function of overhang length for template strand overhang (blue) or non-template strand overhang (green) containing DNA substrates. Error bars represent S.E. (D) Schematic of FRET-pair labelled primed DNA substrate comprised of an 18-mer dsDNA and a 5’ 40-mer poly-dT overhang to assess interactions of Mfd with ssDNA. (E-H) Two example FRET trajectories (top panel) and temporal heat maps (bottom panel) are presented for (E) primed DNA alone (n = 100 molecules), (F) Mfd in the presence of ATPɣS (n = 94 molecules), (G) Mfd in the presence of 1 mM ATP (n = 203 molecules), and (H) Mfd(R953A) in the presence of 1 mM ATP (n = 105 molecules).

#### Primed DNA template (Figure 4)

Primed DNA template for smFRET and cryo-EM was first annealed by hybridizing bio_AS18_Cy5 and Cy3_dT40_S18 in 10 mM Tris/HCl 7.6, 50 mM NaCl, 5 mM MgCl_2_ at a final concentration of 10 µM of Cy3_dT40_S18 and 1.05-fold excess of the bio_AS18_Cy5 complementary strand. DNA was diluted to 50 pM in 200 µL of 1 × TB prior to immobilization in the flow cell (Figure 4).

### 637 nm transcript lifetime experiments

Lifetime of the Cy5 dye was measured by directly exciting *in situ* assembled ECs with a 637 nm laser illumination with a flux of 4 W cm^-2^ (140 mW Vortran, Sacramento, CA, USA) with 200 ms time resolution. Emissions were collected using a dichroic mirror (Dio1-R405/488/561/635, Semrock, Rochester, NY, USA). 100−150 molecules were imaged for each condition. Images were corrected for non-homogeneous laser illumination, and the fluorescence of the molecules (corrected for background) in the entire field of view was calculated on a per pixel basis as a function of time using Fiji, and is presented as mean and standard deviation in Figure 3.

A single exponential decay process best described the observed decay of the signal for the EC alone, but insufficiently described the decay in case of ECs incubated with Mfd and ATP. Therefore, we proceeded with globally fitting the data with a sum of two exponential decay processes. Global fitting was performed by fitting the signals to equation (1) for the two conditions (subscript 1 ≡ EC alone, and 2 ≡ EC incubated with Mfd and ATP):

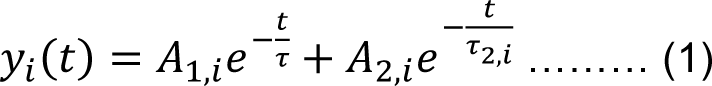

while specifying *A*_2,1_ = 0 and τ_2,1_ = 0 for the signal measured from the EC alone. Here, τ represents the photobleaching rate, and τ_2,i_ describes additional processes influencing the lifetime of the Cy5. Global fitting was performed in OriginPro 2018 using the Levenberg-Marquardt iteration algorithm and iterations were terminated when Chi-square reached a tolerance of 10^-9^ signifying convergence. Optimized values are for EC alone (A_1,1_ = 0.973 ± 0.005; τ = 11.3 ± 0.1 s, A_2,1_ = 0) and for EC incubated with Mfd and ATP (A_1,2_ = 0.49 ± 0.02, τ = 11.3 ± 0.1 s; A_2,2_ = 0.52 ± 0.02, τ_2,2_ = 4.6 ± 0.2 s).

### smFRET Data Analysis

Single-molecule data acquisition was carried out using NIS Elements software. Then, single-molecule intensity − time trajectories and FRET − time trajectories were generated in MASH-FRET software (https://rna-fretools.github.io/MASH-FRET). Heat maps represent aggregated FRET signal for FRET-ting molecules detected in the experiment. Overlaid contour plots represent population deciles, *i.e,* the smallest contour represents the top 10^th^ percentile of the population, the next largest contains the 20^th^ percentile and so on. The projection on the time axis represents the number of observations over time, and the projection on the FRET axis represents the ensemble FRET signal. Transition density plots (TDPs) represent transitions between states fit to individual, idealized trajectories using vbFRET (http://vbfret.sourceforge.net/). Plotted here are kernel density plots of the 2D histograms of the data representing transitions between consecutive states. Data analysis and figures are plotted using custom written Python scripts provided in the supplemental methods.

### Surface plasmon resonance

#### Activation of SPR chip

SPR experiments used a BIAcore T200 instrument (Cytiva) using a streptavidin (SA) coated sensor chip to study the binding kinetics of Mfd to reconstituted ECs, or various DNA substrates. All experiments were carried out at 20 °C with a flow rate of 5 μL min^-1^, unless specified otherwise. The SA chip was activated with three sequential 1 min injections of 1 M NaCl, 50 mM NaOH, then stabilized by 1 min treatment with 1 M MgCl_2_.

#### Immobilization of 49-mer dsDNA substrates (Supplemental Figure 2)

The 49-mer dsDNA substrate was prepared by hybridizing NT_49_bio (final concentration of 1 μM) with two-fold excess of TS_49 (Supplemental Table 1) in 1 × TB in a final volume of 120 μL by heating in a heat block at 90 °C for 5 min followed by slow cooling until room temperature was reached. NT_49_bio:TS_49 dsDNA substrate was resuspended in 1 × SPR buffer (30 mM Tris/HCl, pH 7.6, 40 mM KCl, 5 mM MgCl_2_, 0.005% (*v*/*v*) surfactant P20, 0.25 mM EDTA, 0.5 mM dithiothreitol) to a final concentration of 2.5 nM (of the biotinylated oligo) and introduced onto the SPR chip for immobilization, followed by a wash step with 1M KCl. The signal from the dsDNA template corresponded to 106 response units (RU). Measurement of the Mfd binding affinity for the immobilized dsDNA ligand was performed as described below.

#### Immobilization of reconstituted EC on biotinylated 49-mer dsDNA (Supplemental Figure 2)

The RNAP:template (TS_49):RNA(RNA_15) complex was formed using the procedure described for *in situ* reconstitution of ECs for smFRET measurements. Following this, biotinylated non-template strand (NT49_bio; 10 μL of 10 μM working stock in water) was added to the mixture, which was incubated at room temperature for 15 min. Next, 50 μL of nickel NTA resin suspension were washed and equilibrated with 1 × TB. This resin resuspended in 50 μL of 1 × TB was then added to the mixture containing the RNAP, template:RNA hybrid and non-template strand followed by incubation at 37 °C for 15 min to allow binding of His-tagged RNAP complexes to the resin. The remaining steps were then performed at room temperature. First, supernatant from this suspension was isolated by centrifugation on a mini-benchtop centrifuge for 10 s. The pelleted resin was washed with 150 μL of 1 × TB supplemented with 1 M KCl for 2 min. This suspension was then subjected to centrifugation and the supernatant was discarded. Following this, 150 μL of elution buffer (1 × TB + 500 mM imidazole) were then applied to the resin followed by gentle mixing by pipetting. Finally, the suspension was centrifuged and the eluate containing reconstituted ECs was collected. All reactions containing RNAP were performed in LoBind microcentrifuge tubes (Eppendorf).

The eluate was then applied to a size exclusion column (Wyatt SEC analytical column 030S5) previously equilibrated with 30 mM Tris/HCl, pH 7.6, 150 mM KCl, 15% (*v*/*v*) glycerol, 0.5 mM EDTA, 1 mM dithiothreitol to separate biotinylated ECs from RNAP alone and collected in 200 μL fractions. Fraction 7 (F7) was found to contain the EC as verified by smFRET studies. Immobilization of these reconstituted ECs was performed by resuspending 40 μL of F7 in 200 μL of SPR buffer and introduction into the flow cell. Immobilization was stopped when the signal reached 1380 RU over the background. This was followed by treatment with 1M KCl (10 μL min^−1^) that served to displace incorrectly assembled or non-specifically bound RNAP from the chip surface. The resulting signal corresponded to 550 RU over the background. ECs thus immobilized were found to be stable for several days. Measurement of Mfd binding affinity for the immobilized EC ligand was performed as described below.

#### Immobilization of primed DNA (Supplemental Figure 4)

Primed DNA templates were assembled *in situ* (on the chip surface) in two stages. First, the biotinylated strand was immobilized in three of the four channels in the chip. Single-stranded biotinylated DNA substrates (bio_dT40_S18, bio_AS18, and bio_AS18_Cy5; Supplemental Table 1) were diluted to 1 nM in SPR buffer and immobilized in channels 2, 3, and 4 respectively. This corresponded to 150 RUs of bio_dT40_S18 (channel 2), 75 RU of bio_AS18 (channel 3), and 80 RU of bio_AS18_Cy5 (channel 4). Next, the complementary non-biotinylated strand (2 μM in 1 × SPR buffer) was introduced into the flow cell to hybridize the template *in situ* at 5 μL min^−1^. This led to a further increase in signal corresponding to 39 RU (channel 2, complementary strand: AS18), 138 RU (channel 3, complementary strand: dT40_S18) and 115 RU (channel 4, complementary strand: Cy3_dT40_S18) upon complete hybridization. Following this step, the flow cell was washed with 1 M MgCl_2_. Measurement of Mfd binding affinity to immobilized primed DNA template was performed as described below.

Single-stranded biotinylated DNA substrate (bio_AS18) was diluted to 1 nM in SPR buffer and immobilized (∼48 RU) onto the flow cell of the SA chip by flowing at 5 μL min^−1^. This was followed by flowing in the complementary non-biotinylated strand (5 μM in 1 × SPR buffer) with defined lengths of the ssDNA (dT)_0,3..15_ overhang until the signal reached saturation indicating complete hybridization (typically corresponding to a comparable increase in RU). Following this step, the flow cell was washed with 1 M MgCl_2_. Measurement of Mfd binding affinity was performed as described below.

#### Titration of Mfd (Supplemental Figures 2 and 4)

Following the immobilization of the DNA substrates or the EC, in each case, binding studies were performed by injecting increasing amounts of Mfd in the indicated concentration range in the SPR buffer supplemented with 0.5 mM ATPɣS. For measurements of Mfd binding to dsDNA (Supplemental Figure 2: NT_49_bio: TS_49), reconstituted EC (Supplemental Figure 2) and primed DNA substrates, an optimized concentration range of [0, 10, 20, 40, 80, 160, 320, 640] nM of Mfd was used. For measurements of Mfd binding to ssDNA substrate (bio_dT40_S18), an optimized concentration range of [0, 0.25, 0.5, 1, 2, 4, 8, 16] μM was used. Two conditioning steps of flushing the flow cell with 300 mM KCl for 30 s at 10 µL min^−1^ were carried out prior to injection of Mfd. Mfd-ATPɣS was flowed in for 150 s in the association phase, and dissociation was observed for 200 s in 1 × SPR buffer lacking any Mfd or nucleotide cofactor.

### Analysis of sensorgrams

Sensorgrams were subtracted from the signal from unmodified flow cell (flow channel 1), then zero-subtracted (injection at 0 μM Mfd or mutant) and *R*_eq_ values, generated by averaging response values in the sensorgram’s steady-state region from the appropriate Mfd concentration range, fit against [Mfd] using a 1:1 steady state affinity (SSA) model incorporated in the BIAevaluation software 4.0.1 (Cytiva):

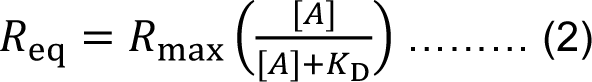

where *R*_max_ corresponds to the response when all the immobilized DNA ligands on the surface are saturated with the analyte *A*, *K*_D_ is the dissociation constant, and [*A*] is the concentration of analyte (Mfd or mutant) in solution. In general, raw sensorgrams and fit curves (where applicable) were exported using Excel, then plotted in Origin, and final Figures prepared in Adobe Illustrator CC.

### Electron microscopy sample preparation, data acquisition and analysis

Mfd stock (40 μM; 30 μL) was dialyzed overnight in 1 L of 1 × TB at 4 °C in a Slida-A-lyzer mini dialysis unit with 3,000 MWCO (Thermo Fisher) to reduce the amount of glycerol in the sample. Following this, protein was recovered and concentrated to a 1.67 mg mL^−1^. For the sample with non-hydrolyzable ATP abalog ADP-AlF_x_, to the mixture of Mfd (7.8 μL at 1.6 mg mL^-1^) and fluorescent oligonucleotides (10 μL of 10 μM stock), ADP, NaF and AlCl_3_ were added to final concentrations of 2, 5 and 0.5 mM, respectively. This reaction mixture was incubated at 37°C for 5 min to allow binding of Mfd to the template. The samples were clarified by centrifugation at 20,000 × g for 10 min at 4 °C before grid preparation. Samples (3 µL) were applied to glow-discharged Quantifoil R 1.2/1.3, Au 400 grids. The grids were blotted at 6 °C for 3.5 s at 100% humidity with no extra blot force in an FEI Vitrobot Mark IV and plunged into liquid ethane. The grids were stored in liquid nitrogen until use.

Data were collected using Thermo Fisher Talos Arctica microscope at 200 kV equipped with a Falcon III (Thermo Fisher) camera in electron counting mode. Micrographs were acquired at a nominal defocus range of −1.0 to −1.9 μm, with a pixel size of 0.74 Å and a total dose of 50 e^-^ Å^−2^ accumulated over 50 s and fractionated across 50 movie frames.

Data were processed in cryoSPARC v2 (46). For the sample with ADP-AlF_x_, 472 images were manually selected after motion correction and CTF estimation for final processing. A total of 110562 particles were selected after 2D classification and subjected to 3D classification in *ab initio* reconstruction followed by heterogeneous refinement using four *ab initio* classes. Particles (31718) from class 3 were used for 3D refinement, which resulted in a 3D reconstruction at 5.2 Å resolution (0.143 FSC gold standard; Supplemental Figure 5A-C).

#### Model fitting

Molecular dynamics with explicit solvent was used to fit the model of Mfd on primer DNA into the cryo-EM density map (47). The starting model was comprised of the Mfd motor domain (PDB ID 2EYQ) with idealized B-DNA with the same sequence as used for the cryo-EM experiments but with a dT_12_ overhang instead of dT_40_. The system was equilibrated for 10 ns. The resulting model was then used as the basis for modeling the complex in COOT (48) with minimal distortions to protein secondary structure and DNA. The model was then subjected to restrained positional refinement in phenix real_space_refine (49). The final model is comprised of 382 residues from Mfd and 47 nucleotide residues. The model was visualized in Chimera (50).

#### Measurement of 2AP fluorescence

Double-stranded (ds) and bubble substrates used in 2AP experiments were hybridized at a concentration of 20 μM in 1 × TB by heating to 95 °C on a heat block for 5 minutes followed by turning off the heat source and allowing the reaction to cool down to room temperature overnight. dT6_S18_v2_11_2AP or dT6_S18_v2_12_2AP were hybridized with either AS18_v2_2AP or AS18_v2_2AP_bubble (sequences are provided in Supplemental Table 2) to create four substrates: (a) 11_2AP_ds ≡ dT6_S18_v2_11_2AP: AS18_v2_2AP (b) 11_2AP_bubble ≡ dT6_S18_v2_11_2AP: AS18_v2_2AP_bubble (c) 12_2AP_ds ≡ dT6_S18_v2_12_2AP: AS18_v2_2AP and (d) 12_2AP_bubble ≡ dT6_S18_v2_12_2AP: AS18_v2_2AP_bubble. Starting with a stock concentration of 40 μM of Mfd and 20 μM of DNA substrates, protein and DNA were diluted 10-fold into 1 × TB in a total sample volume of 120 μL and 100 mM ATPɣS stock was supplemented to a final concentration of 0.5 mM as appropriate. For purposes of background subtraction, we collected emission spectra of Mfd (4 μM) alone, Mfd (4 μM) and ATPɣS (0.5 mM), or ds or bubble substrates alone (2 μM). All samples were incubated for at least 20 minutes at room temperature (24 °C) prior to measurements.

Spectra were collected on a Cary Eclipse in FP8300 emission mode. Samples were deposited in a clean quartz cuvette with a total sample volume of 120 μL. Samples were excited at 330 nm (slit width 5 nm) and emission spectra were collected in the range from 350 nm – 460 nm with a slit width of 5 nm and a scan rate of 30 nm/min. The averaging time for each measurement was set as 1 s and data were collected at an interval of 0.5 nm. The PMT voltage was set to 800.

Two independent repeats were collected for each of the ds and bubble substrates (11_2AP_ds, 11_2AP_bubble, 12_2AP_ds, 12_2AP_bubble). Each spectrum was blanked for buffer, and the difference spectrum was calculated by subtracting the blanked spectrum of the ds substrate from that of the bubble substrate. Fluorescence enhancement was calculated by subtracting the signal measured in the presence of Mfd and ATPɣS from the signal measured in the presence of Mfd alone for both substrates. The fluorescence emission of either ds or bubble DNA substrate (2 μM) was measured in the presence of Mfd (4 μM) alone or with 0.5 mM ATPɣS and blanked by subtracting the corresponding emission spectrum of a sample of either Mfd (4 μM) or with 0.5 mM ATPɣS in 1 × TB, and lacking the DNA substrate. Three repeats were performed for 11_2AP_ds, 11_2AP_bubble and 12_2AP_bubble and two repeats were performed for the 12_2AP_ds substrate. The average difference spectrum was then calculated at each wavelength in the emission spectrum and a five point moving average was further used to smooth this signal for purposes of visualization. In the figures, individual corrected measurements are plotted as points, and the average is plotted as a line.

## RESULTS

### Rationale for a smFRET assay to study transcription elongation

RNAP in the elongation phase can exist as a backtracked complex (potentially extensively) where the 3’ end of the transcript is extruded into the secondary channel (2), a pre-translocated conformation prior to nucleotide addition, and a post-translocated conformation following nucleotide incorporation. We monitored occupancy of these states during transcription elongation by assembling functional ECs on dye-labeled, fully-complementary transcription bubbles *in situ* by adopting a previously described strategy to study RNA Pol II dynamics (51) in combination with well-established approaches to reconstitute functional *E. coli* ECs (45). Specifically, we introduced a Cy3 donor dye on the upstream DNA and a Cy5 acceptor on the RNA (Figure 1A; Supplemental Figure 1B) at sites that are predicted to exhibit high smFRET signals within the dynamic range of the technique (Supplemental Note 1). Three features of this strategy enable precise dissection of the dynamics of molecular mechanisms of Mfd-dependent remodeling of the EC: (i) the use of synthetic RNA and DNA oligonucleotides enables site-specific introduction of dyes in the phosphodiester backbone that serve as FRET pairs, and consequently, the ability to probe conformational changes between various parts of this complex; (ii) smFRET affords sub-second temporal resolution (200 ms in our experiments) enabling reliable measurements of dynamic interactions on the timescale of seconds to minutes, and (iii) the ability to specify RNA sequences that cannot hybridize with upstream DNA limits extensive backtracking of ECs. Therefore the observed population is expected to reside in the pre- and post-translocated registers.

### An analysis workflow to monitor transcription elongation

Functional RNAP ECs were assembled *in situ* on the microscope stage at room temperature (24 °C) by flowing a solution of transcription buffer containing RNAP that had been pre-incubated with the template:RNA hybrid into a microfluidic device with surface immobilized biotinylated non-template DNA (Figure 1A). After complex assembly, the flow cell was washed with a high-ionic strength solution (transcription buffer supplemented with 1M KCl) followed by rapid exchange with a buffer solution supporting transcription but without ribonucleotides (NTPs). These steps enabled the retention of only completely reconstituted ECs that each contain a Cy3-labeled template and a Cy5-labeled transcript (see Supplemental Table 1 for sequences). Excitation of these ECs with 532 nm laser illumination revealed a population of molecules exhibiting FRET. The smFRET signal from these surface immobilized molecules was collected in individual movies (acquisitions) lasting 120 s, with a temporal resolution of 200 ms. The smFRET signals were analysed in five ways: 1. Regions of interest containing individual molecules were used to generate trajectories by plotting the evolution of the background-corrected smFRET signal over time. 2. The aggregated trajectories were visualized in a ‘temporal heat map’ to monitor the progress of the reaction at the ensemble level (below). 3. The trajectories were idealized and transitions between various kinetic states were visualized at the ensemble level using ‘transition density plots’ (below). 4. Individual trajectories were further parsed into various classes depending on the types of transitions between FRET states observed in the idealized trajectories. 5. Finally, having access to the residence time of molecules in individual states, we quantified lifetimes of the kinetic intermediates observed in specified FRET regimes (below). The overall analysis workflow is described below.

First, examination of smFRET trajectories describing the smFRET signal between the Cy3 donor and the Cy5 acceptor (see Methods) revealed that the assembled complexes resided in stable high FRET states centered at 0.9 (Figure 1B top panel, upper trace; Supplemental Figure 1C for example donor-acceptor intensity traces). In addition, a small fraction (14% of 318 trajectories) also exhibited mid-FRET states centered at 0.6 (Figure 1B top panel, lower trace).

Next, these states were recapitulated in temporal heat maps (Figure 1B, middle panel). Heat maps are scatterplots describing how the smFRET signal evolves in time for the ensemble population. The heat maps represent the raw data, and the projections on the Y-axis represent the ensemble FRET measurements (of all FRET trajectories), while the projection on the X-axis displays the evolution over time of all FRET signals of the entire population of molecules with a time resolution of 0.2 s. The loss of signal over time is attributable to the photobleaching of individual dyes during the course of the acquisition due to continuous laser illumination (Figure 1B, middle panel). Here, the overlaid black curves reflect contours parsing the data into deciles.

Next, we plotted transition density plots (TDPs) to obtain an ensemble overview of the reaction. Features in TDPs correspond to interconversions between individual states observed in individual trajectories. Since the majority of the complexes thus assembled exhibited time-FRET trajectories that fluctuate around 0.9 FRET, and a minority exhibited transitions to the 0.6 FRET state, we pursued an unbiased approach to visualize transitions between various FRET states. FRET trajectories were idealized (see Methods) and the detected states were plotted in TDPs (Supplemental Figure 1D, ‘EC alone’) to visualize interconversion among different states and the directionality of those changes (52). In this representation, features that are only present in one half of the plot on either side of the diagonal manifest as asymmetric features and denote irreversible transitions between states on the timescale of the observation. As demonstrated by the TDP, the interconversions between the states were reversible (based on the displayed symmetry of the FRET populations across the diagonal), and were restricted to the high FRET regime. Within the paradigm that FRET changes reflect conformational changes, a straightforward interpretation of these data is that the complex undergoes relatively minor, but reversible, conformational changes.

To gain insight into the reaction at the level of individual trajectories, we assigned smFRET signals between 0.7–1 as being ‘high’, 0.4–0.7 as ‘mid’ and 0–0.4 as being ‘low’. Trajectories were then sorted as ‘static’ (black bars; bottom panel Figure 1B) if the signal resided in exactly one state, and dynamic otherwise. Further, dynamic trajectories were parsed into ‘high’, ‘mid’ or ‘low’ categories if all the states visited by the molecule were within the ranges defined above. Molecules exhibiting transitions between two adjacent regimes were classified as ‘high–mid’ or ‘mid–low’ as appropriate, and molecules exhibiting transitions among all three regimes were counted in the ‘high–low’ class. The lower panels represent column plots of the percentages of the trajectories in each class, and the total number of molecules observed is indicated in the top right in each panel. Here, for ‘EC alone’ 65% of the ECs exhibited static FRET signals. 10% exhibited dynamics confined to the high-FRET regime, and 14% and 9% of observations respectively exhibited transitions in the high–mid and high–low FRET regimes.

Finally, with access to the idealized trajectories, we extracted lifetimes of ECs under various conditions in specified FRET regimes (for example: 0.35–0.5 FRET). The residence time of individual states in this regime were then fit to single exponential (denoting a single Poisson process with a single rate-limiting step) or Gamma functions (denoting the existence of multiple steps with comparable lifetimes). This kinetic analysis enabled us to better understand the nature of the intermediates underlying the measured FRET signals. For example, in the case of the EC alone, we recovered a lifetime of 5.0 ± 0.8 s from a single-exponential fit indicating the existence of a single intermediate with a residence time of ∼5 seconds in the mid-FRET regime (Supplemental Figure 1). Together, the various analyses serve to provide an overview of static and dynamic behavior by the individual molecules in the data set.

#### Monitoring transcription elongation by RNAP

To assign states, and to test that the complexes are catalytically functional, we challenged complexes with the full set of deoxyNTPs (dNTPs), all ‘wrong’ NTPs (defined as the full set of NTPs lacking the next cognate NTP which is GTP in this case; Figure 1A) and the complete set of NTPs. In this assay, transcript elongation is expected to increase the distance between the FRET pair dictated by the geometry of the advancing EC, leading to a drop in FRET. Since dNTPs are selected against incorporation into the transcript (53), we expected that smFRET signal from the reconstituted ECs would not drop. Consistent with this expectation, complexes exposed to dNTPs revealed a high-FRET population (Figure 1C). Compared to the behaviour of ECs in the absence of any nucleotides, a greater fraction of the population exhibited static molecules (Figure 1C 74%, compared to 65% in the absence of any dNTPs, Figure 1B). Within the dynamic population, ECs mainly exhibited dynamics in the ‘high-mid’ (10%), and the ‘high’ (6%) FRET regimes, confirmed by the observation of strong peaks in these regimes in the TDPs (Supplemental Figure 1D, ‘EC, dNTPs’). Thus, dNTPs shift the equilibrium such that ECs reside primarily in the high-FRET regime, populated by catalytically inactive complexes (see below).

In contrast, non-complementary NTPs dynamically shifted the equilibrium at the population level between molecules occupying the the high- and mid-FRET range, to a population exhibiting FRET signals mainly in the mid-FRET range (Figure 1D). Fewer static trajectories were observed in this case (40% compared to 65% in the case of ‘EC alone’ column plot in Fig 1B), and dynamic trajectories exhibited reversible transitions in all three FRET regimes (Figure 1D, column plots). This is further supported by the observations in the TDP that the interconverting populations exhibited symmetric peaks across the diagonal indicating that the ternary complex visits the observed states reversibly (Supplemental Figure 1D, ‘EC, wrong NTPs’). Notably, strong peaks are observed in the mid-FRET regime in the TDP (Supplemental Figure 1D, ‘EC, wrong NTPs’) at the expense of the transitions in high-FRET regime seen in the case of ECs alone or supplemented with dNTPs (Supplemental Figure 1D, ‘EC alone’ and ‘EC, dNTPs’). The lack of asymmetric transitions in the TDP when ECs were incubated with all wrong NTPs/dNTPs indicates either that mis-incorporation of the wrong nucleotide does not occur appreciably during the observation of the ECs, or that if it occurs, it is rapidly reversed.

‘Walking’ the complex by supplying a subset of the ‘correct’ complementary NTPs revealed gradual conversion from the high-FRET to a mid-FRET population (compare projections on the Y-axes in the heatmaps in Supplemental Figure 1E−G). When presented with the complete set of NTPs, the EC exhibited unidirectional conversion of the FRET signal from high- to low-FRET values (Figure 1E). Notably, the TDP (Supplemental Figure 1D), and indeed the individual traces, often revealed a conversion of the high-FRET states to the low-FRET states *via* the mid-FRET states. 30% of trajectories in the data set (Figure 1E column plot) exhibited dynamics from the high to the low FRET regime. In this class of trajectories, 30% of the ECs exhibited a single direct transition, 57% exhibited a step-wise transition to the low-FRET regime by visiting the mid-FRET regime (intermediate; Supplemental Figure 1D, ‘EC, all NTPs’) and the remainder (13%) exhibited high-mid dynamic trajectories prior to a single-step transition from a high-FRET state to a low-FRET state. It is likely that trajectories exhibiting a single-step transition conceal an intermediate step with a lifetime below our time resolution.

To better characterize the mid-FRET states observed in these experiments, we plotted distributions of lifetimes of individual states in each condition (Supplemental Table 2, Supplemental Figure 1H and Supplemental Note 2). Here, the EC exhibited a lifetime of 5.0 ± 0.8 s, and this did not change appreciably in the presence of dNTPs (5.7 ± 1.5 s) or correct NTPs (5.5 ± 1.3 s) although, a slightly longer lifetime of 7.8 ± 0.9 s was detected in the presence of the wrong NTPs. In conjunction with the TDPs, we infer that despite comparable lifetimes, the exit from the mid-FRET states is favored toward entry into high-FRET states for all conditions except in the presence of the correct NTPs consistent with a functional complex.

Collectively, the data indicate that the complexes assembled *in situ* represent functional ECs that exist in a catalytically ‘inactive’ kinetic state and a catalytically ‘poised’ kinetic state. The observations that complexes often first transition to the mid-FRET state prior to addition of NTPs suggests that this represents a catalytically poised kinetic state. We tentatively assigned the high-FRET states centered at 0.9 to a catalytically inactive kinetic state. While drawing one-to-one correspondence between structural and kinetic states is an exercise that should be approached cautiously, the smFRET signals measured in these complexes can illuminate the potential structural states underlying the kinetic states observed in our experiments. Estimates of the expected FRET efficiency based on the distances between the Cy3-Cy5 dye pair for the pre- and post-translocated conformations of the EC are in good agreement with the measured FRET efficiencies measured in our experiments (Supplemental Figure 1B and Supplemental Note 1). Therefore, a reasonable interpretation of these experiments is that the EC reversibly transits between the pre- and post-translocated structural states that manifest as catalytically inactive and catalytically poised kinetic states. Consistent with this framework, these experiments indicate that the EC is highly dynamic in the presence of incorrect NTPs, but exhibits irreversible forward translocation only in the presence of the correct NTPs consistent with catalysis and transcript elongation.

### NusG promotes residence of ECs in the catalytically poised state

Next, we tested this assignment of states by assessing the response of the complex to the elongation factor NusG (Figure 2A), which is predicted to retain ECs in the catalytically poised state. Addition of NusG indeed resulted in an increased percentage of dynamic ECs in high-mid and high-low FRET regimes (together 33% of the population in Figure 2B, compared to 23% for ‘EC alone’ Figure 1B; column plots) accompanied by a reduction of molecules in the high-FRET regime (2% in Figure 2B compared to 10% in Figure 1B; column plots). Individual trajectories revealed a conversion from the high-FRET state into the mid-FRET state and fluctuation around the mid-FRET state (Figure 2B), consistent with our assignment of the mid-FRET as the catalytically poised state. We did not detect evidence of protein induced fluorescence enhancement, suggesting the lack of interactions between Cy3 and NusG bound to the upstream edge of the transcription bubble.

Incubation with all wrong NTPs (UTP, ATP, CTP) in the presence of NusG revealed a greater percentage of dynamic ECs exhibiting transitions in the high-low and mid-low FRET regimes compared to NusG alone (together 28% for ‘EC, NusG wrong NTPs’ in Figure 2C compared to 21% for ‘EC, NusG’ in Figure 2B; column plots) at the expense of fewer transitions in the high-mid regimes. Reversible short-lived transitions to low-FRET states were also evident in individual trajectories (Figure 2C) and in the TDP (Supplemental Figure 2A). The lifetime of the EC in the presence of NusG and all wrong NTPs in the mid-FRET state (6.9 ± 0.9 s) was similar to that of the EC alone incubated with all wrong NTPs (7.8 ± 0.9 s) while being longer than that of the EC incubated with NusG alone (4.2 ± 0.4 s) in the absence of any nucleotide (Supplemental Figure 2B).

Importantly, the complete set of NTPs revealed efficient, irreversible conversion to the low-FRET regime (0 – 0.4) from the mid-FRET regime (heatmap, Figure 2D and Supplemental Figure 2A) in the presence of NusG. Individual trajectories exhibited unidirectional conversion (Figure 2D, top panel). Only 23% of the population accounted for static molecules exhibiting only a single state, almost three-fold fewer than observed in the case of ECs incubated with NusG alone or with the wrong NTPs (Figure 2B and 2C, column plots). Of the remaining, 69% of molecules exhibited dynamics in the high-low and mid-low FRET regimes. In this case, the TDPs exhibited additional asymmetric features (Supplemental Figure 2, ‘EC, NusG all NTPs’). The lifetime of the EC in the presence of NusG and all correct nucleotides was comparable to the lifetime of the EC in the absence of any nucleotide or in the presence of correct nucleotides (Supplemental Table 2, Supplemental Figures 2B, 1H and Supplemental Note 2). The asymmetric features reflect processive extension of the transcript through nucleotide incorporation and associated forward translocation by RNAP. Together, these observations are consistent with the role of NusG as a suppressor of backtracking of RNAP (7–9), indicating that NusG modulates the equilibrium between the catalytically inactive and active kinetic states by plausibly promoting residence of the complex in the post-translocated register.

### Nucleotide-bound Mfd translocates the EC forward

Next, we assessed the interactions of Mfd with ECs, armed with the assignment of states in the high-FRET regime as catalytically inactive states (consistent with the pre-translocation register), and those in the mid-FRET states as a catalytically poised states (consistent with the post-translocation register). To characterize the binding of Mfd to the EC, we first measured the affinity of Mfd for pre-assembled ECs or the dsDNA alone using surface plasmon resonance (SPR). Since Mfd interacts poorly with DNA in bulk assays in the absence or presence of ATP, measurements of binding of purified Mfd to either DNA or the EC were carried out in the presence of slowly hydrolyzable ATPɣS. SPR measurements revealed a dissociation constant (*K*_D_) of 170 ± 20 nM (Supplemental Figure 2C) for 49-mer dsDNA alone, consistent with measurements by other groups (11, 24). Binding to reconstituted ECs on a 49-mer exhibited similar affinity with a *K*_D_ of 127 ± 6 nM (Supplemental Figure 2C). These data are consistent with the model that dsDNA serves as a major binding partner for Mfd-ATPɣS in the context of the EC as previously demonstrated (12,13,22,24). It is notable that whereas association curves of Mfd to dsDNA exhibit fast saturation, binding of Mfd to RNAP ECs does not reach full saturation on the timescale of the association phase. This may be attributable to the presence of two types of binding substrates, for example, non-specific binding to dsDNA alone, or specific binding to the EC.

We then investigated how interactions of Mfd with RNAP influence the conformational dynamics of the transcription bubble using our smFRET assay (Figure 3A). Consistent with the lack of stable binding of Mfd with DNA in the absence of ATP, only 49% of trajectories of ECs exhibited observable transitions in the smFRET measurements (*n_obs_* = 77, Figure 3B bottom panel). However, this is greater than the number of dynamic trajectories observed in the case of ECs alone (35%, Figure 1B, column plots). Notably, compared to the EC in the absence of Mfd (Figure 1B), a greater percentage of dynamic trajectories exhibited transitions in the high-mid FRET regime in the presence of Mfd (compare Figure 1B column plot ‘EC alone’ high-mid class: 14% and Figure 3B column plot ‘EC, Mfd’ high-mid class: 24%). These trajectories exhibiting reversible transitions from the high- to mid-FRET states even in the absence of any nucleotide co-factor reflect failed attempts at stable engagement of Mfd with the EC.

Incubation with Mfd-ATPɣS resulted in approximately the same fraction of dynamic trajectories of ECs visiting additional mid-FRET states as in the case of Mfd alone (Figure 3B and 3C, Supplemental Figure 3A ‘EC, Mfd-ATPɣS’; column plots). Additionally, the loss of molecules exhibiting dynamic trajectories in the ‘high-mid’ and ‘high’ FRET regimes was compensated by an increase in the number of static trajectories, and molecules exhibiting dynamics in the ‘high-low’ and ‘mid’ FRET regimes (column plots in Figures 2C and 2B respectively). Curiously, in the presence of Mfd-ATPɣS, the EC exhibits apparently quantized transitions in the transition density plot compared to incubation with Mfd alone (Supplemental Figure 3A ‘EC, Mfd’ *vs.* ‘EC, Mfd-ATPɣS’). The presence of additional low-FRET states in the transition density plot (Supplemental Figure 3A ‘EC, Mfd-ATPɣS’) indicates that binding of Mfd-ATPɣS displaces the EC forward. ATPɣS alone did not account for these observations (Supplemental Figure 3B). Together the data reveal that Mfd can isomerize the EC merely upon binding to the nucleotide-starved EC. The efficiency of this isomerization is appreciably greater in the presence of ATPɣS.

### Nucleotide-bound Mfd activates the EC

Having established that Mfd-ATPɣS can promote forward translocation of the EC, we investigated whether this binding is sufficient to activate the EC. To test whether ECs pre-incubated with ATPɣS for 10 minutes could further incorporate rNTPs, we introduced GTP and UTP into the flow cell and measured the FRET signal (ATP was left out as it can be utilized by Mfd). As expected, molecules exhibited a rapid conversion to low-FRET states evidenced in individual trajectories, as well as in the ensemble heat map (Figure 3D). The characteristic lifetime of ECs in the intermediate states was recovered from a gamma distribution to be 0.7 ± 0.2 s (Supplemental Figure 3C), almost eight-fold lower than the mean lifetime of the intermediate state for ECs incubated with NTPs (5.5 ± 0.9 s) recovered from a single-exponential distribution. This striking, almost eight-fold reduction in the lifetime of the intermediate formed by Mfd-ATPɣS provides evidence that nucleotide-bound Mfd primes the EC for transcription elongation. Considering that the EC is dynamic on the template in the presence of nucleotides (Figure 1), a plausible physical basis for this observation is that Mfd-ATPɣS bound to the upstream edge of the transcription bubble acts as a ratchet that restricts diffusion to the pre-translocated register (backtracking), and thus reduces retention of the RNAP in non-productive registers on the template.

To quantify the ability of ECs to incorporate nucleotides in the absence or presence of Mfd-ATPɣS, we compared histograms of the ensemble FRET states detected in the idealized trajectories for the two conditions (Figure 3E). Fitting the histogram to a sum of Gaussians revealed three populations centered at FRET values of 0.91, 0.63 and 0.09 for the EC incubated with GTP, UTP and ATP. We assigned these as the EC alone, catalytically poised EC, and ECs that have extended the transcript following NTP incorporation, respectively. Similarly, we recovered three populations centered at FRET values of 0.93, 0.7 and 0.18 for ECs that were pre-incubated with Mfd-ATPɣS followed by incubation with GTP and UTP. We assigned these three populations as EC alone, ECs acted upon by Mfd-ATPɣS and ECs that have incorporated two NTPs.

In each case, the recovery of the population at 0.9 FRET indicates the presence of catalytically inactive ECs (consistent with the pre-translocated register). Further, the recovery of the two populations with mean FRET at 0.63 (95% confidence intervals, lower bound (LB): 0.48, upper bound (UB): 0.77) and 0.7 (LB: 0.65, UB: 0.75) suggests that the FRET signal of the catalytically poised state (consistent with the post-translocated register) formed by the EC in the presence of NTPs is indistinguishable from that of the intermediate formed by the EC in the presence of Mfd-ATPɣS. Importantly, a third population emerges in time (Figure 3E) when NTPs are present. This population exhibited a FRET state of 0.09 (LB: 0.05, UB: 0.12) when the EC was incubated with GTP, UTP and ATP. In comparison, for ECs incubated with Mfd-ATPɣS and GTP and UTP, this population exhibited a FRET signal of 0.18 (LB: 0.15, UB: 0.2). The difference in the mean FRET values (0.09 *vs*. 0.18) likely reflects the ability of ECs incubated with GTP, UTP and ATP to synthesize a longer transcript compared to ECs incubated with only GTP and UTP.

Comparison of the fraction of the population in the low FRET states corresponding to nucleotide incorporation by the EC alone or the EC pre-incubated with Mfd-ATPɣS (Figure 3E) revealed a two-fold greater efficiency of incorporation after pre-incubation with Mfd-ATPɣS: 35% of all FRET states observed in the reaction corresponded to low-FRET states when ECs were pre-incubated with Mfd-ATPɣS *vs.* 15% in the absence of Mfd-ATPɣS (Supplemental Note 3). Together the data enable us to conclude that nucleotide-bound Mfd serves to promote forward translocation of the EC into a catalytically poised kinetic state (consistent with the post-translocated register) that is competent for transcription elongation when presented with NTPs. This finding is surprising because previous work has suggested that ATP hydrolysis dependent translocation on the upstream DNA is necessary for productive loading of Mfd on to the EC (12) and transcription rescue (13). Hence, our data (Figure 3C-E) reveal an efficient alternative loading pathway of nucleotide-bound Mfd that yields activated ECs without the need for extensive ATP hydrolysis on upstream DNA. This efficient loading and activation is clearly observed on our DNA template where backtracking is prohibited; presumably, ATP hydrolysis-dependent engagement of Mfd plays a role in rescuing ECs from extensively backtracked states.

### A single rate limiting gating intermediate determines transcription termination

Next we probed the effect of Mfd in the presence of ATP on the fate of the EC. Incubation of *in situ* assembled ECs with Mfd and ATP (1 mM) revealed a greater number of molecules exhibiting dynamics (Figure 3F, 58%, compared to 44% in the presence of ATPɣS and 49% in the absence of nucleotide) across all three regimes including molecules that exhibited efficient conversion to the mid- and low-FRET regimes (Figure 3F, heat map and trajectories). To further explore the nature of these transitions, we visualized them in a TDP (Supplemental Figure 3A, ‘EC, Mfd-ATP’), which revealed a dynamic population in the mid-FRET regime. Measurement of the lifetime of the EC incubated with Mfd-ATP in this intermediate state revealed a lifetime of 6.2 ± 0.7 s, comparable to that of the EC incubated with Mfd alone (7.1 ± 2 s) or in the presence of Mfd-ATPɣS (6.1 ± 1.4 s) (Supplemental Table 2, Supplemental Figure 3C and Supplemental Note 2). Strikingly, the lifetime of the EC in this mid-FRET state did not exhibit a dependence on the concentration of ATP in the range from 10 µM to 10 mM suggesting that stoichiometric amounts of ATP are sufficient to promote conversion to low-FRET states (Supplemental Table 2 and Supplemental Figure 3C). We note that ATP alone was not responsible for the conversion of the EC to low-FRET states in the absence of Mfd (Supplemental Figure 3D).

Compared to the wild-type, the Mfd(R953A) mutant (17), known to be proficient in ATP hydrolysis while deficient in RNAP displacement and transcription termination exhibited inefficient conversion of the EC to the low-FRET states in the presence of ATP (Figure 3G). Notably, unlike wild-type Mfd, a greater number of complexes were found to be static (compare panels 3F and 3G 42% *vs*. 63%) and residing mainly in the high-FRET regime (Figure 3G, heatmap, projection on Y-axis). Dynamic trajectories exhibiting high-mid, high-low and mid-low (Figure 3G, column plot, together 33%) transitions were less frequent compared to wild-type Mfd (Figure 3F column plot, together 55% respectively). TDPs revealed a greater fraction of individual transitions in the high- to mid-FRET regime for Mfd(R953A) (Supplemental Figure 3A ‘EC, Mfd(R953A)-ATP’ *cf.* Supplemental Figure 3A ‘EC, Mfd-ATP’ panel). The lifetime of ECs incubated with Mfd(R953A) and ATP in the mid-FRET regime was identical to that of ECs incubated with Mfd alone in the absence of nucleotide (Supplemental Figure 3C). Intriguingly, ECs incubated with Mfd(R953A)-ATP exhibited a pattern of quantized transitions that is similar to that of ECs incubated with Mfd-ATPɣS in the transition density plots (Supplemental Figure 3A, compare ‘EC, Mfd-ATPɣS’ and ‘EC, Mfd(R953A)-ATP’), suggesting that the mutation in this residue serves to reveal distinct aspects of Mfd’s binding to the EC.

Together, the data indicate that both wild-type Mfd and translocase deficient Mfd rescue RNAP from the catalytically inactive states accessible in our substrate to catalytically-poised states, albeit with varying efficiencies. However, only the wild-type Mfd can further process the EC in the presence of ATP efficiently. The ECs exhibiting low-FRET states in our system represent complexes in which the distance between the donor–acceptor pair is outside the dynamic range of efficient FRET as might be expected when the RNAP is forward translocated well past the stall site, and/or when the Cy5-labeled RNA transcript is lost. The observation that the EC fails to return to the mid- or high-FRET states in the presence of ATP (Figure 3F) indicates the occurrence of an irreversible reaction, most consistent with an irreversible loss of RNA observed during transcription termination (54). In this model, Mfd(R953A) fails to displace the transcript from the EC, consistent with prior findings (17).

### Loss of transcript marks exit from the gating intermediate

To confirm that the exit from the mid-FRET regime observed in the ECs incubated with Mfd-ATP corresponds to loss of the transcript and not photobleaching of the Cy5 dye, we measured the residence time of the Cy5-labeled RNA through direct excitation using a 637 nm laser (see Methods). Here, the distribution of the lifetimes of the Cy5 signal in samples containing ECs alone on the surface revealed a single species with a lifetime of 11.3 ± 0.1 s corresponding to the photobleaching lifetime of Cy5 due to direct excitation by the laser (Figure 3H). On the other hand, ECs incubated with Mfd in the presence of ATP (1 mM) displayed a lifetime distribution that was best fit by two exponentials: a short-lived species with a lifetime of 4.6 ± 0.2 s (amplitude: 52 % of the population) in addition to the longer-lived species detected previously, corresponding to the photobleaching of Cy5 with a mean lifetime of 11.3 ± 0.1 s (Figure 3H). This short-lived species exhibits a lifetime that matches well the lifetime of the catalytically poised EC incubated with Mfd-ATP measured using smFRET (4.6-6.2 s; Supplemental Figure 3C). Further, the fraction of dynamic trajectories observed in the FRET experiments (Figure 3F column plot, 58%) agreed well with the fraction of the population losing the RNA transcript (52%). Thus, the data demonstrate that exit from this intermediate indeed corresponds to loss of the RNA transcript. In essence, we have uncovered a single rate-determining intermediate that serves as a ‘molecular timer’ counting down the time beyond which elongation is no longer an option.

### Motivation for a new model for Mfd-mediated transcription termination

Structures of Mfd bound to dsDNA and ECs containing non-complementary transcription bubbles have revealed a series of translocation intermediates that are formed during the loading phase of Mfd on to the EC (12, 22). On the basis of these structures a model has been proposed in which Mfd-mediated transcription termination may occur in a step-wise rewinding of the transcription bubble upon the advancement of Mfd translocating on the upstream DNA (Model I). Our smFRET experiments aimed at uncovering the mechanism of transcription termination revealed the presence of a single intermediate (Figure 3F) that is formed upon binding of nucleotide-bound Mfd to the EC. Further, in the presence of ATP (Figure 3D), we detected the exit from this intermediate in a single step corresponding to loss of the RNA transcript indicating transcription termination. Critically, the concentration of ATP did not influence the lifetime of this intermediate (Supplemental Figure 3C). With experimental access to a faster temporal resolution (0.2 s) and an assay that is exquisitely sensitive to the architecture of the transcription bubble, we detected lifetimes that are distributed according to a single exponential function, indicating the presence of a single, rate-determining step with a lifetime of ∼6 s. In this intermediate, the smFRET signal remains constant, indicating that the complex does not undergo dynamic conformational changes. At the end of this lifetime, we found that the RNA transcript is ejected indicating completion of transcription termination. Therefore, these results do not provide experimental support for a model that hypothesizes step-wise transcription termination where Mfd rezips the transcription bubble one nucleotide at a time through processive ATP hydrolysis on this timescale. However, we note that the lifetimes of rezipping intermediates would have to be significantly less than our temporal resolution, on the order of milliseconds, and these intermediates must occur at the end of the lifetime of the intermediate detected here.

### Mfd specifically recognizes template-strand ssDNA

Hence, we turned our attention to the other leading model (Model II) for forward translocation proposed on the basis of structures of the eukaryotic homologs of Mfd (32, 41). In this model, TRCF melts and extends the transcription bubble on the upstream edge, and makes direct contacts with either the template or non-template DNA through key residues. We set out to identify whether Mfd makes contacts with ssDNA in substrates mimicking the upstream edge of the transcription bubble by employing a combination of approaches: First, we assessed whether Mfd exhibits a preference for binding to template or non-template ssDNA overhangs at junctions, second, we employed smFRET experiments to detect whether and how Mfd contacts template ssDNA, and finally, we visualized Mfd bound to a primed DNA template using cryo-EM.

To investigate whether Mfd can recognize ssDNA in substrates mimicking the upstream edge of the transcription bubble, we measured the DNA binding affinity of Mfd to 18-mer dsDNA substrates containing either template or non-template strand overhangs (Figure 4A, B; Supplemental Table 1). First, we interrogated whether length of the non-template strand overhang influenced the affinity of Mfd for the DNA substrate (Figure 4A–C). We measured the dissociation constant of Mfd binding to dsDNA (18-mer) with varying lengths of non-template strand overhangs (dT_x_ in which x = 0, 3, 6, 9, 12, or 15 nucleotides; Figure 4B). In the presence of slowly-hydrolyzable ATPɣS, the dissociation constant did not exhibit a dependence on the overhang length (Figure 4C, green trace) indicating that Mfd is insensitive to the presence and length of non-template strand overhangs. Next, we measured binding affinity of Mfd-ATPɣS to dsDNA substrates (18-mer) with varying lengths of unpaired 5’ template strand overhangs (dT_x_ in which x = 0, 3, 6, 9, 12, or 15 nucleotides; Figure 4A–C). In this case, the binding curves revealed an inverse relationship between the length of the ssDNA overhang and the strength of interaction: the longer the overhang, the weaker the binding. *K_D_* values increased from 90 ± 4 nM (for 18-mer ds DNA alone) to 220 ± 10 nM (for 18-mer dsDNA with a 15 nt overhang). Finally, Mfd bound a 58-mer ssDNA with a *K_D_* of ∼4 µM (Supplemental Figure 4A) which is 20-fold weaker than binding to dsDNA alone, consistent with prior estimates (55). These data establish the ability of Mfd to distinguish between template and non-template strand overhangs.

In this assay, the Mfd(R953A) mutant similarly exhibited a loss of affinity as a function of template strand overhang length up to a dT6 overhang in the presence of ATPɣS (Figure 4C, dark blue trace). The dissociation constant for binding a substrate containing a 6 nt template-strand overhang was 2.3-fold weaker than that for wild-type Mfd. Substrates containing overhangs larger than 6 nt were bound markedly more strongly, and approached the binding affinity of Mfd for these substrates (Figure 4C). These data reveal that only contacts made by Mfd on the template ssDNA overhang serve to antagonize its binding to dsDNA, establishing that the non-hydrolyzable ATP analog-bound conformation of Mfd exhibits strand specificity at DNA forks. Further, the results highlight a curious role of R953 in mediating these contacts on the template strand and a dependence on the length of the overhang. We note that these data are not adequately explained by available structural models that reveal the R953 contacts only the non-template strand in the Mfd-transcription complex (12).

To further understand these ATP-dependent contacts of Mfd with ssDNA we employed a previously described smFRET assay that has been used to study translocase activity (56). We used a DNA substrate comprised of an 18-mer dsDNA duplex (mimicking upstream DNA) and a 5’ dT_40_ overhang mimicking the template DNA (Figure 4D) with a FRET pair positioned on either end of the ssDNA overhang (Supplemental Table 1). Mfd binds this substrate with a *K*_D_ of 390 ± 10 nM (see Supplemental Note 4 and Supplemental Figure 4B−D). In smFRET experiments, this substrate exhibited an average FRET of 0.33 ± 0.15 (Figure 4E) in the absence of Mfd, reflecting rapid sampling of various conformations accessed by the entropic coil formed by ssDNA (Figure 4E and Supplemental Figure 4E ‘template overhang DNA’).

In the presence of ATPɣS, the smFRET signal from the Mfd-bound substrate revealed additional distinct populations at higher FRET values of 0.6 and 0.8 (Figure 4F and Supplemental Figure 4E ‘Mfd-ATPɣS’). Transitions among these states were further enhanced in the presence of ATP as the substrate exhibited a continuous spectrum of mid- and high-FRET states (Figure 4G and Supplemental Figure 4E ‘Mfd-ATP’), in addition to the distinct states detected in the presence of ATPɣS. In this case, the higher FRET states were only transiently accessed indicating that ATP hydrolysis plays a role in exiting from these high−FRET states to low−FRET states. This observation is consistent with a picture in which Mfd-ATP compacts the ssDNA to bring the dyes close to each other, resulting in a high FRET signal, and subsequent ATP hydrolysis induces a lower-affinity conformation in which Mfd-ADP binds the ssDNA overhang very weakly resulting in an abrupt return to the baseline FRET value of 0.33. This ‘compacting’ behaviour is in stark contrast to the processive ‘reeling’ behaviour of PcrA and related helicases on this substrate (56).

To verify that the dynamics observed in this assay reflect a dependence on ATP hydrolysis, we studied binding to this substrate by the ATP hydrolysis-deficient but DNA-binding competent E730Q mutant of Mfd (27, 57); this Mfd(E730Q)-bound template exhibited high-FRET states extremely rarely compared to the wild-type protein (Supplemental Figure 4F, 4G). Therefore, the observation that Mfd(E730Q)-ATP is unable to stably engage with template strand ssDNA, unlike Mfd-ATPɣS suggests that the Mfd(E730Q)-ATP and Mfd-ATPɣS pre-hydrolytic states are sufficiently different that only the wild-type Mfd retains the ability to compact ssDNA efficiently in the ATP-bound state.

Unexpectedly, the R953A mutant did not exhibit high FRET states in the presence of ATP, indicating that the ability to hydrolyze ATP alone is insufficient to compact ssDNA on primed DNA templates (Figure 4H). Therefore, we conclude that both, the ability to hydrolyze bound ATP and the presence of an intact R953 are essential for Mfd to engage the 5’ template strand ssDNA overhang. We propose that R953 serves to ‘hook’ template strand ssDNA, and ATP binding serves to ‘pull’ it closer to the junction resulting in the high FRET signal detected in this assay with wild-type Mfd. In turn, ATP hydrolysis leads to exit from the high-FRET state reflecting slippage of the ssDNA from the motor domain. We note that the recent structures of Mfd translocating on dsDNA (12) indicate that R953 along with key residues in the translocation motif (TRG: translocation in RecG (14)) of Mfd, specifically R929 and Q963, contacts the non-template strand in dsDNA. From these structures it is not immediately obvious how the template strand overhang is contacted by Mfd in the smFRET assays. The results presented here uncover a new role for R953 in making critical contacts with template ssDNA distinct from its previously described role in translocation on dsDNA. To reconcile the previous observations with our findings, we propose that residue R953 contacts the non-template strand during the initial loading phase (12) and switches to a contact with the 5’ template strand ssDNA during the remodelling phase. This finding explains the deficiency of this mutant of Mfd in coupling ATP hydrolysis to removal of RNAP.

Intriguingly, each of the three Mfd variants also exhibited a distinct sub-population of molecules centered at ∼0.1 FRET. Here, whereas ATPɣS promoted the formation of this state almost immediately, Mfd (or mutants) in the presence of ATP entered this state in a time window ∼5–20 s after introduction into the flow cell (Figure 4F-H and Supplemental Figure 4H). The presence of the low-FRET peak and the excursions to high-FRET states suggest that multiple independent binding modes may underlie the FRET signal observed here.

### Mfd melts dsDNA in primed-DNA substrates

To further understand how Mfd binds template strand overhang−containing DNA, we visualized Mfd bound to the dye-labeled smFRET substrate by cryo-EM (Figure 5A, Supplemental Table 1). Full-length Mfd bound to the 5’ tailed substrate in the presence of transition-state analog ADP-AlF_x_ was subjected to single particle cryo-EM analysis (Supplemental Figure 5A) and a ∼5.2 Å resolution map was obtained (real resolution ∼8 Å, Supplemental Figure 5B, C). This reconstruction yielded EM density that could at best accommodate the motor domains of Mfd (residues 575−956 representing domains 5 and 6); other domains are presumably flexible and could not be seen in the EM density maps. In addition the reconstruction could accommodate 18 bp of dsDNA and 11 nt of the 5’ dT (template strand) overhang (Figure 5B).

**Figure 5:**
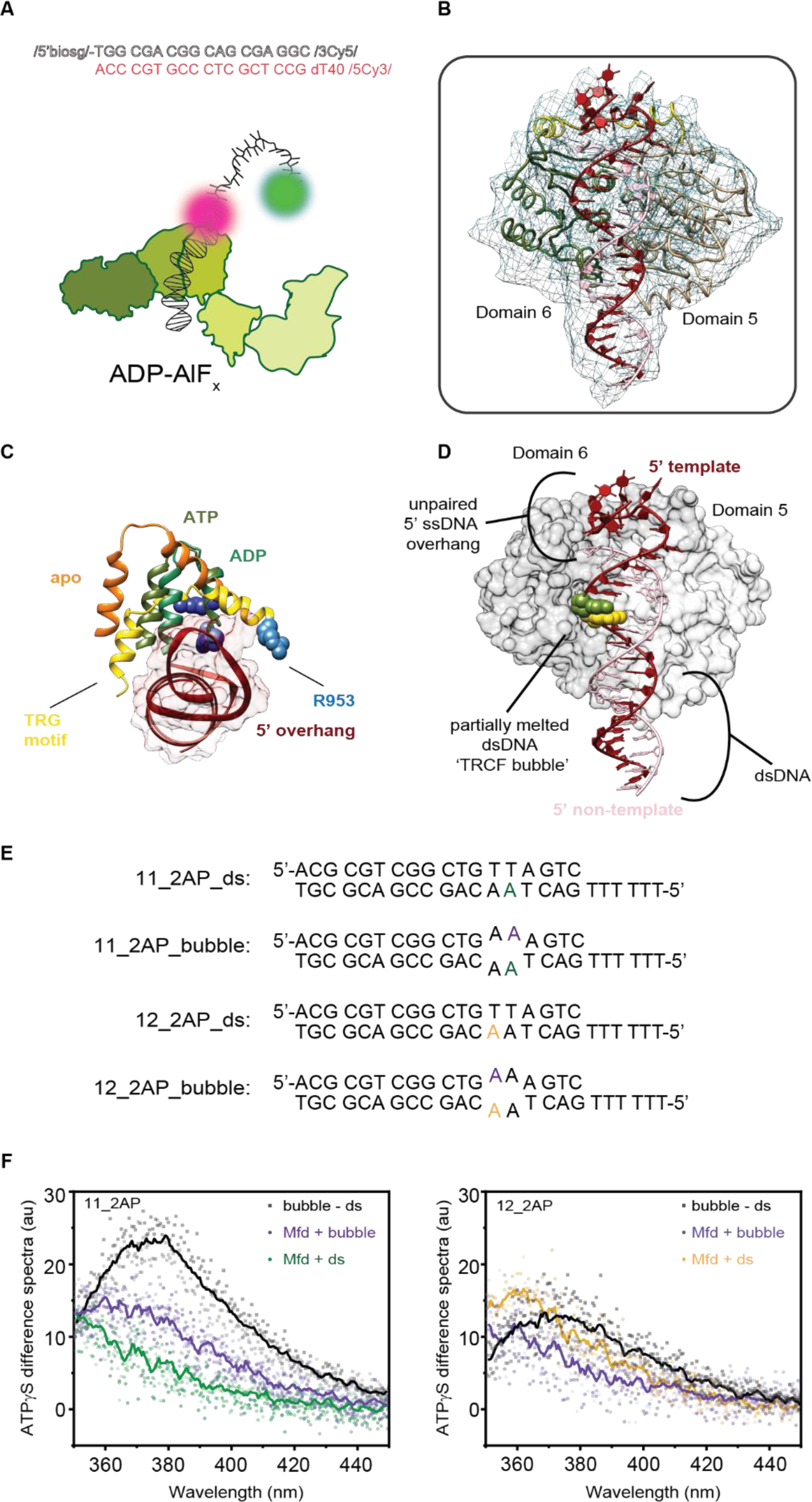
Interactions between Mfd and primed DNA. (A) Sequence of primed DNA substrate and schematic of complex subjected to cryo-EM. (B) Front view of the molecular model derived from the cryo-EM reconstruction showing Mfd (residues 575-956) including domains 5 (tan) and, 6 (olive), template (dark red) and non-template (pink) DNA. DNA (template: residues 28−58; non-template: 1−18) are shown in ribbon representation to illustrate the path of the DNA. (C) Comparisons of the conformations of the TRG motif in apo (orange; PDB 2EQY); ADP-bound (green, PDB 6X4W); ATP-bound (olive; PDB 6X43) and on primed-DNA (yellow, ADP.AlFx) in structures aligned to the motor domain in PDB 6X43. R953 is shown as blue spheres for the ATP, ADP and apo structures. The primed DNA substrate is shown in ribbon and surface representation. (D) The model suggests dsDNA near the junction is melted to form the TRCF bubble. Position 11 from the 5’ end of the template is shown in green, and position 12 is shown in yellow on the template strand. (E) Sequences and schematic of DNA substrates used in Mfd-dependent DNA melting experiments. (F) Difference spectra are shown for the fluorescence of 2AP in bubble substrates compared to the respective ds substrates in the absence of Mfd or ATPɣS (black). Difference spectra showing increase in fluorescence upon addition of ATPɣS to a sample containing Mfd and either the bubble substrate (purple) or ds substrate (panels show data for substrate containing 2AP at position 11(left; green in panel D) or 12 (right; yellow in panel D)). Lines indicate five-point moving averages, and individual squares/circles indicate data from 2-3 independent repeats.

The cryo-EM map reconstructed here suggests the dsDNA on the upstream side of the motor domains and the placement of the motor domains with respect to dsDNA to be consistent with previously reported complexes (11, 12) (Figure 5B). However, EM density for residues 917 onward, containing the TRG motif, appeared to be absent from the side of domains 5 and 6, and the EM density of the DNA as it passes from the upstream to the downstream side becomes inconsistent with dsDNA (Figure 5B). Using molecular dynamics flexible fitting (47) to equilibrate the structure in density suggests that the TRG motif has dropped into the path of the dsDNA (yellow ribbon, Figure 5C), resulting in the partial melting of the duplex with only 13 residues remaining double stranded. The motif forms an arch that interacts with the template DNA (Figure 5C). There are precedents for conformational plasticity in the TRG motif: the pair of helices forming the bulk of the TRG motif are almost orthogonal in the apo-structure of Mfd (PDB ID 2EYQ; Figure 5C, orange) with the sulfonate group from 4-(2-hydroxyethyl)-1-piperazine ethanesulfonic acid bound between the helices, and are parallel in the complex with ATP- and ADP-bound conformations on upstream dsDNA (PDB ID 6X43 and 6X4W). These movements are attributable to both to shifts in domains 5 and 6 relative to each other and to movements of the helices relative to each other. The conformation of domains 5 and 6 suggested here may represent a mode of engagement whereby Mfd grabs and stabilizes the template DNA in the upstream edge of the transcription bubble, supporting the inferences drawn from the biophysical investigations.

The molecular model derived from the cryo-EM investigations presents two aspects of Mfd binding that can be experimentally validated. Despite the poor resolution of the cryo-EM data which limits confident assignment of the position of R953, the model is consistent with and supports the observation from the smFRET studies that R953 mediates compaction of the 5’ ssDNA overhang (Figure 4). Additionally, the model suggests that Mfd partially melts about five nucleotides of the dsDNA near the junction forming a ‘TRCF bubble’ (Figure 5D). An additional attempt to obtain an improved higher resolution reconstruction with an unmodified dT_6_ template strand overhang-containing 18-mer ds template yielded a low-resolution reconstruction of limited utility (data not shown). Considering these limitations, we used an orthogonal approach to test the model prediction that binding of Mfd in the presence of a non-hydrolyzable ATP analog-leads to melting of the dsDNA in template strand-overhang containing DNA substrates.

2-Aminopurine (2AP), is a fluorescent analog of adenine that has been previously used to study protein-induced local conformational changes in DNA (58, 59). 2AP readily base pairs with dT and its fluorescence is quenched in dsDNA. We reasoned that local melting of DNA could be observed by monitoring the unquenching of the fluorescence signal that occurs when 2AP is exposed to the solvent. To test whether nucleotide-bound Mfd destabilizes primed DNA templates, we designed two sets of substrates (Figure 5E, 18 bp double-stranded with 6 nt of 5’ template strand overhang) – a first set comprised of completely complementary double stranded substrates (‘ds’) with 2AP at either position 11 or 12 (counted from the 5’ end on the template strand of the substrate; positions are highlighted in the model in Figure 5D), and a matched set where the non-template strand contains a non-complementary sequence resulting in a ‘bubble’ opposite the introduced 2AP at position 11 and 12 (Figure 5D and 5E).

First, we measured the difference in the emission spectra between the bubble and ds substrates (2 µM each) for each of the substrates containing 2AP at position 11 or 12 (Figure 5F black trace) upon excitation at 330 nm. This difference spectrum demonstrates that the fluorescence of 2AP in the bubble substrates is greater than that in the double-stranded substrates, consistent with the expectation that 2AP is quenched in dsDNA. This signal reflects the maximum fluorescence enhancement resulting from exposure of 2AP upon local melting of ds DNA to form a bubble.

Next, the DNA substrate (2 µM, either ds or bubble) was incubated with Mfd (4 µM) in the presence or absence of ATPɣS (0.5 mM), and emission spectra were measured at steady state. Spectra were baseline corrected for Mfd (4 µM) and ATPɣS (0.5 mM) separately and the resulting corrected spectra are plotted in Figure 5F. Incubation of the 2AP bubble substrates with Mfd and ATPɣS resulted in a fluorescence enhancement (Figure 5F, purple trace) for both substrates. Importantly, in the presence of Mfd and ATPɣS, both 2AP-containing dsDNA substrates exhibited an enhanced fluorescence (Figure 5F, green and yellow traces). The increase in fluorescence was greatest when 2AP was introduced at position 12, and was indistinguishable from the maximum fluorescence enhancement on this substrate.

Additionally, we noted that the emission peak is blue shifted in the presence of Mfd and ATPɣS compared to samples containing the DNA substrates alone (black). Blue shifts in the emission spectrum of 2AP reflect a change in the local environment to one possessing a lower dielectric constant (59, 60), suggesting a hydrophobic interaction with residues of Mfd. Together, the data provide independent confirmation of the prediction from the cryo-EM derived molecular model that Mfd melts DNA substrates containing a template-strand overhang in the presence of non-hydrolyzable ATP analogs, and maintains contact with the melted DNA. Consistent with this scenario, Mfd-ATPɣS binds a 49-mer dsDNA substrate containing a non-complementary bubble of 9 nucleotides with an affinity of 395 ± 7 nM as assessed by SPR (Supplemental Figure 5D).

Together, the findings converge on a ‘melt, hook and pull’ model to describe how Mfd may contact DNA substrates mimicking the upstream edge of the transcription bubble: SPR experiments reveal that Mfd specifically recognizes the template strand and this binding antagonizes binding to dsDNA. Mutation of R953 weakens this hooking interaction with single-stranded template DNA. Further, the cryo-EM data and 2AP fluorescence experiments suggest that Mfd binds and partially melts the dsDNA near the junction, contacting the unpaired template ssDNA *via* the conserved R953 residue in the TRG motif. Finally, the smFRET experiments reveal that Mfd dynamically cycles between high- and low-affinity states in an ATP-hydrolysis dependent manner on the template ssDNA overhang, pulling and compacting the template strand in this process, in an R953 dependent manner. These findings lay down a framework for a new model of Mfd- and ATP-mediated remodelling of the EC.

## DISCUSSION

In this work, we set out to understand dynamic modulation of EC activity by the action of interacting partners. We detected that NusG promotes the residence of the EC in catalytically poised states. We established that a ‘molecular timer’ intermediate is formed by Mfd, and ATP-hydrolysis-dependent exit from this state determines the fate of the EC. Our attempts to understand this intermediate resulted in the discovery of a novel DNA-binding mode in which Mfd binds and melts upstream fork mimic DNA substrates and contacts template-strand ssDNA *via* a hook residue. Together, the results provide insight into the transcription termination reaction and enable us to propose a comprehensive model describing how events unfold after initial engagement of the EC with the transcription-repair coupling factor.

Interactions of Mfd with the EC are complex and proceed *via* three stages: recognition, loading and remodelling. In the first stage, Mfd recognizes a paused/ stalled EC and is recruited (13,23,24,27,61) through a conserved RNAP-interacting domain (13) (Figure 6A). Mfd can recognize extensively backtracked (24), reversibly paused as well as stalled complexes (61). In the second stage, EC-tethered Mfd undergoes a series of concerted conformational changes (20) that enable it to translocate on upstream dsDNA and ‘load’ onto the EC (12, 22) (Figure 6B). In the third stage, Mfd either reactivates paused transcription (13, 24) or terminates transcription (16, 23). Alongside these activities, Mfd recruits its binding partners UvrA_2_B to the site during transcription-coupled DNA repair (16,26–29).

**Figure 6:**
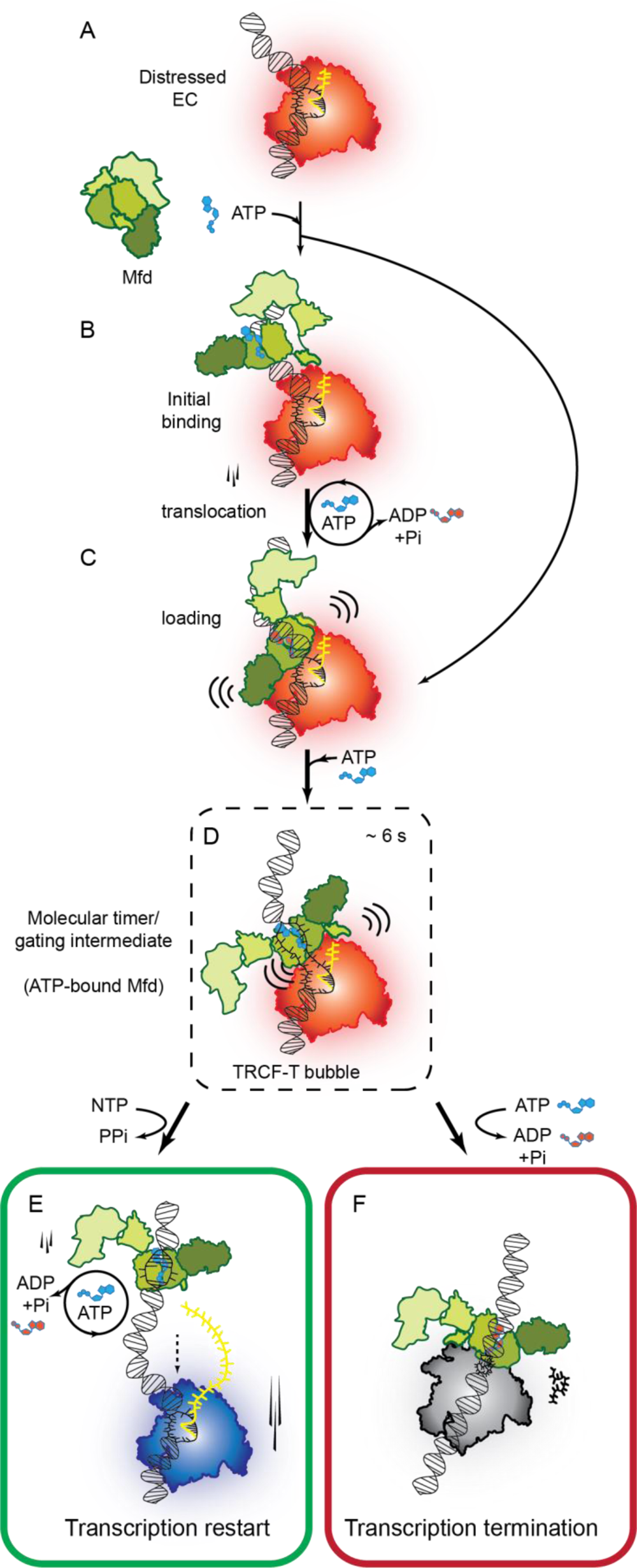
Model for Mfd-mediated transcription modulation. (A) Mfd interacts with the EC weakly in the absence of ATP, however, (B, C) ATP binding and hydrolysis enables Mfd to bind and translocate on the upstream DNA as described in ref(12) resulting in loading of Mfd on to the EC. In an alternate loading pathway detected here, nucleotide-bound Mfd may also load on to the EC without the need for ATP-hydrolysis. (D) Nucleotide-bound Mfd rescues the EC from catalytically inactive state, into a molecular timer intermediate in the post-translocated register with a lifetime of ∼4−6 s. In this ATP-bound state, we propose that Mfd melts the upstream DNA to form an extended transcription-repair coupling factor – transcription (“TRCF-T”) bubble. Further, in this state R953 contacts the template ssDNA in the TRCF-T bubble. Melting of this upstream DNA is sufficient to promote residence of the EC in the post-translocated register. (E) In the presence of NTPs on an undamaged template, transcription can be resumed explaining the basis of the proposed ‘release and catch-up’ model(24). (F) On damaged templates or under conditions of nucleotide starvation or in the presence of impediments to the progress of the EC, the bound ATP is hydrolyzed to promote transcription termination accompanied by the loss of the RNA transcript.

While the initial stages of recognition and loading are relatively well understood, precisely how remodelling of the transcription complex proceeds is still unclear. Our study sheds light on hitherto mysterious aspects of Mfd-RNAP interactions central to the remodelling phase and presents a new model for resolution of a critical intermediate. The smFRET-based elongation modulation assay revealed that Mfd can engage the EC in an ATP independent manner, albeit infrequently, and manifest transient, reversible conformational changes in the EC. Stable association requires ATP or at minimum non-hydrolyzable ATP, likely required to impart a dsDNA binding conformation in the motor domain. A recent study reported ATP-hydrolysis dependent translocation intermediates formed during initial recognition of nucleotide-starved ECs leading to loading of Mfd on the EC (12). Here we found that Mfd loads efficiently on to the EC to yield active complexes even in the absence of ATP hydrolysis on substrates on which backtracking is forbidden.

Following a period of dynamic remodeling in which the EC transits from high- to mid-FRET states, Mfd-ATP, Mfd-ATPɣS and Mfd(R953A)-ATP each convert the EC into a single long-lived gating intermediate (Figures 3 and 6D) hovering in the mid-FRET range. This intermediate controls the transition to low-FRET states. Efficient entry into this state is granted to ECs incubated with Mfd in the presence of ATP or ATPɣS, whereas ECs incubated with Mfd(R953A)-ATP enter this state inefficiently. The observation that Mfd-ATPɣS is sufficient to promote entry into this state indicates that binding of ATP alone is a sufficient requirement for the formation of the gating intermediate.

The suggestion from the cryo-EM-derived molecular model that Mfd-ADP-AlF_x_ might bind and melt primed DNA templates to unwind dsDNA (Figure 5), and partial confirmation of this model in the 2AP fluorescence experiments provides insight into this transition. Only 13 nucleotides of the 18-mer DNA template are consistent with being dsDNA, with the remainder being partially melted. The lack of observations of dsDNA melting on templates lacking the single-stranded overhang in a recently reported structure (11) suggests that Mfd requires access to unpaired ssDNA *via* critical residues in the TRG motif to melt the dsDNA.

We propose that in the context of the upstream dsDNA, ATP- or non-hydrolyzable analog-bound Mfd partially melts about five nucleotides of dsDNA to produce a bubble, henceforth referred to as the ‘TRCF bubble’ (Figure 5). Considering the three nucleotide proximity of the binding site of Mfd to the upstream edge of the transcription bubble (12), the upstream DNA may further melt allowing the TRCF bubble to fuse with the transcription bubble to produce the isomeric TRCF-transcription (‘TRCF-T’) bubble (Figure 6D). Since prior experiments did not detect bubble formation by Mfd on dsDNA lacking an overhang (23), a more likely possibility is that Mfd merely extends the transcription bubble by further melting the upstream edge, resulting in essentially a TRCF-T bubble. While our study was under review, an independent study reported that ‘a highly negatively supercoiled domain forms between Mfd and RNAP’, consistent with our report of a TRCF-T bubble (62) suggesting further support for a *bona fide* ability of Mfd to melt DNA.

We note that available structures of Mfd-[ATP or analog]-EC do not report melting of the upstream DNA or a TRCF-T bubble (12, 22), but this is not surprising as these structures containing intact transcription bubbles reflect loading, and not remodelling intermediates. While a structure of the Mfd-EC bound to the TRCF-T bubble signifying a remodelling intermediate is not as yet forthcoming, precisely such an intermediate has been observed in the Rad26-Pol II structure where an extended TRCF-T bubble of 17 nucleotides is observed (32). Similarly, human CSB was also reported to bind large transcription bubbles in a recent study (41).

Our cryo-EM derived model suggests that R953 lines the path of the template overhang (Figure 5). The observation that Mfd(R953A)-ATP fails to stably engage the ssDNA overhang in our smFRET study supports a model in which this residue (potentially along with other residues in the translocation motif) serves to stabilize the fork at the upstream edge of the TRCF bubble, or template strand ssDNA in the TRCF-T bubble. We propose that binding of R953 to the ssDNA template occurs simultaneously with the melting of the upstream dsDNA. Evidently, along with a role in DNA translocation in which R953 contacts the non-template strand, R953 has a newly discovered role in hooking and stabilizing template strand ssDNA on substrates mimicking the upstream edge of the transcription bubble. Remarkably, a similar element responsible for hooking the non-template DNA in the upstream fork in the case of CSB was recently identified in a study published while ours was in revision (41). Here, the authors demonstrated that residue F796 in CSB serves as a hook by inserting itself in the upstream fork in the non-template strand. In the yeast homolog Rad26, K613/R614 in the wedge motif together serve to hook the template strand (32). While the wedge motif itself is absent in Mfd, R953 appears to play a functionally analogous role as a hook in the prokaryotic model. In all three cases, this hook serves to couple ATP binding and hydrolysis to remodeling of the EC. These similarities – the presence of a hooking element and destabilization of the upstream dsDNA – between the prototypical members of this class of transcription-coupled DNA repair factors establish a more profound conservation of mechanism of action.

We propose that the formation of the TRCF-T bubble results in forward translocation of the EC to the post-translocated register. We envision this occurs *via* restriction of diffusion to the pre-translocated register due to physical occlusion by Mfd, and establishment of contacts via R953 accompanied by compaction of single-stranded template DNA. Here, in the presence of the correct NTP and an intact R953, resumption of transcription would lead to faster forward translocation of RNAP, leaving behind Mfd-ATP bound to the TRCF bubble (Figure 6E). In effect this finding explains how Mfd can ‘release’ paused/nucleotide-starved polymerases in the so-called ‘release and catch-up’ model for transcription rescue (24), and how Mfd can discriminate between paused and stalled ECs (23). Indeed, this prediction of our model was verified in the experiment demonstrating that Mfd bound to ATPɣS leads to a significantly truncated residence time in the intermediate state accompanied by loss of FRET consistent with transcription rescue (Figure 3 and Supplementary Figure 3).

To understand what happens once the molecular timer intermediate (Figure 6D) forms, we must first consider that exit from this intermediate is signified by a loss of FRET in the presence of ATP and no other nucleotides. Our measurements of the residence time of the transcript through direct excitation of the Cy5 matched the lifetime of the EC in the intermediate state, indicating that loss of FRET corresponds to loss of the transcript from the complex. This interpretation is supported by previous observations that loss of RNA from the EC occurs rapidly after Mfd binds the EC (25). An important consequence of entry into this gating intermediate is that a finite time window is available beyond which transcription restart is no longer possible. The prolonged residence in the catalytically poised state prior to irreversible remodelling of the EC by Mfd may serve to provide an opportunity to restart transcription at sites of pausing. This observation explains how Mfd can orchestrate transcription restart (24) or termination (23) on distressed polymerases, discriminating paused ECs that can be substrates for transcription restart from irreversibly or DNA damage-stalled ECs that are substrates for transcription termination.

It can be deduced that resolution of this gating intermediate during transcription termination proceeds *via* at least partial, and potentially complete, shrinking of the TRCF-T bubble leaving behind Mfd bound to the TRCF bubble. How might these events play out? Our results demonstrate that the lifetime of this state is only ∼4−6 s at 24 °C. While entry into this state is dependent on the presence of ATP, the observation that this lifetime is independent of ATP concentration in the range from 10 µM to 10 mM suggests that hydrolysis of stoichiometric amounts of ATP, may be sufficient to promote exit from this state. Further, an intact R953 is required for exit from this state. The loss of the transcript signals the resolution of the molecular timer, accompanied by the formation of an intermediate where the EC is extensively remodeled, and the TRCF-T bubble is partially or completely rewound. The cryo-EM derived model presented visualizes how Mfd may compact template DNA in this process.

Mfd recruits the repair machinery to the site of distressed polymerases during transcription-coupled nucleotide excision repair. During global genomic nucleotide excision repair, the damage recognition enzyme UvrA_2_ partially melts putative damage containing dsDNA and loads two copies of the damage verification enzyme UvrB at the site (63). Two UvrB helicases then examine the unpaired ssDNA with single-nucleotide resolution to precisely identify the damaged strand (64, 65). In contrast, during transcription-coupled DNA repair, Mfd recruits UvrA_2_B to the site of the distressed polymerase (28, 29). Compared to global genomic nucleotide excision repair, only a single copy of UvrB is recruited to a site of damage by Mfd through interactions with UvrA_2_. Thus, sequestration of specifically the damage-containing template by Mfd supports the intriguing possibility that Mfd possesses the intrinsic ability to faithfully convey the identity of the damaged strand to complete the handoff to a single UvrB *via* the adaptor UvrA_2_. Further elucidation of transcription-coupled DNA repair intermediates will shed light on this idea.

## AVAILABILITY

All custom scripts used for analysis are available at https://github.com/hghodke/mfd_FRET.

## ACCESSION NUMBERS

CryoEM map has been deposited in the Electron Microscopy Data Bank (https://www.ebi.ac.uk/pdbe/emdb/) under accession code EMD-25408. Coordinates for the EM-derived molecular model have been deposited in the RCSB Protein Data Bank (https://www.rcsb.org/) with accession code 7SSG.

## SUPPLEMENTARY DATA

Supplementary Data are available at NAR online.

## ACKNOWLEDGEMENT

We thank Rachel Mooney and Robert Landick (UW Madison) for generously sharing expression vectors for RNA polymerase and purified NusG used in these studies. We thank Stephen Ralph for use of the fluorometer in the School of Chemistry and Molecular Bioscience.

