## Supplemental Info for "Mechanism of transcription modulation by the transcription-repair coupling factor"

Supplemental Figure 1

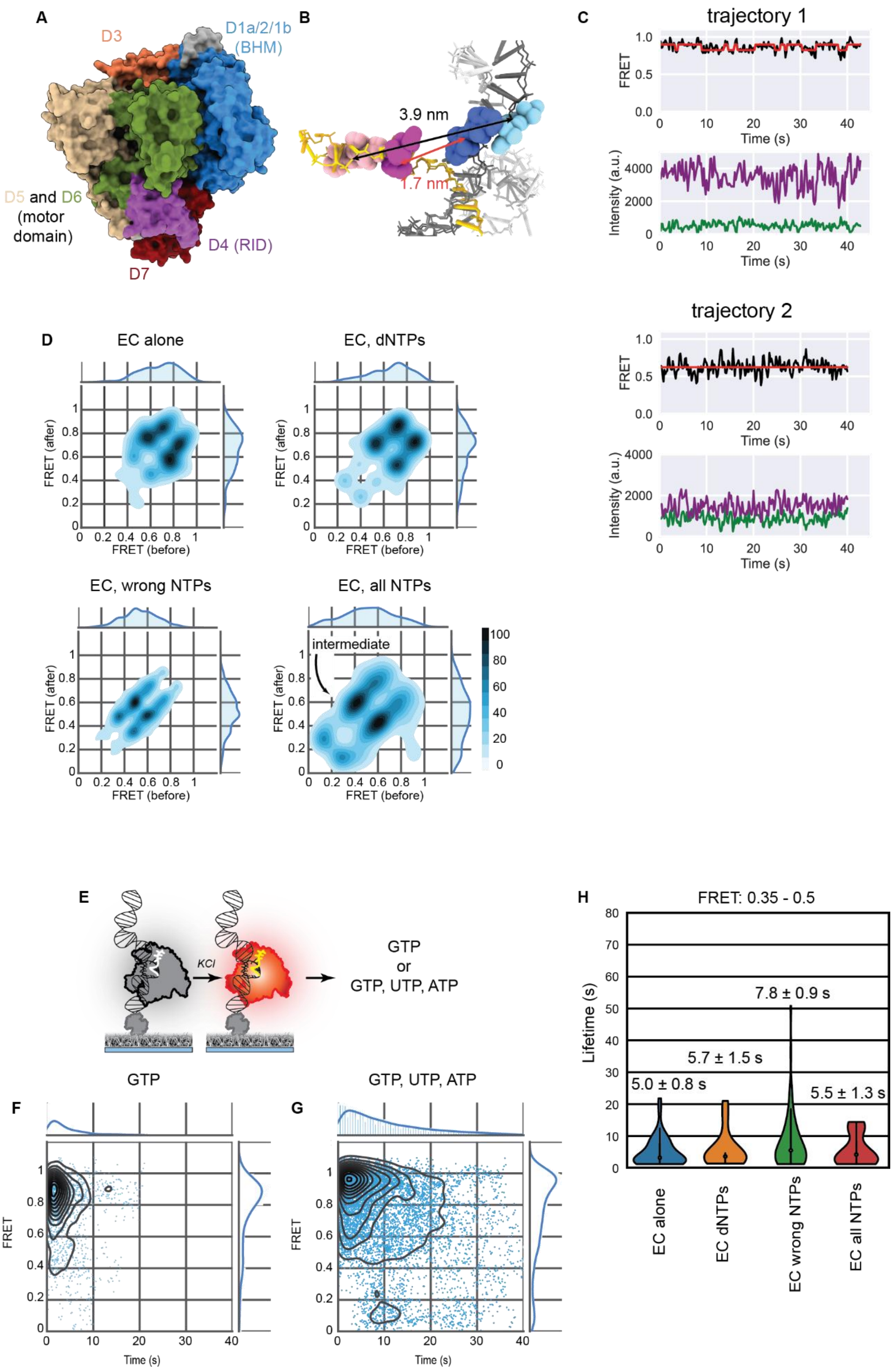

**Supplemental Figure 1: Walking the EC with subsets of correct rNTPs.** ***Related to Figure 1.***

(A) Crystal structure of apo Mfd (PDB 2EYQ) colored by functional modules: UvrB homology module (blue), domain 3 (coral), domain 4 (RNAP interacting domain; purple), motor domains 5 (tan) and 6 (olive), auto-inhibitory domain 7 (dark red).

(B) Schematic of the *E coli* transcription bubble (non-template: light gray, template: dark gray, RNA: gold) with label positions highlighted. Blue spheres represent Cy3 and magenta/pink spheres represent Cy5 inserted in the backbone of the nucleic acids. The distances between the cyanine dyes are indicated for the pre-translocated register (1.7 nm) and post-translocated register (3.9 nm) (see also Supplemental Note 1).

(C) Two example FRET trajectories of the EC alone (as shown in Figure 1B) and corresponding intensity traces for the Cy3 donor signal (green) and Cy5 acceptor (magenta). Trajectories are truncated at the photobleaching step.

(D) Transition density plots for RNAP alone (n = 318 molecules), in the presence of dNTPs (n = 79 molecules), with non-complementary (‘wrong’) NTPs (n = 179 molecules) and the full set of NTPs (n = 81 molecules) respectively.

(E) Schematic of *in situ* assembled FRET-pair labeled ECs incubated with GTP (the next correct NTP) (n = 63 molecules) or with GTP, ATP and UTP (n = 87 molecules). Temporal heat maps of the reaction for experiments conducted in the presence of GTP (F) or GTP, ATP and UTP (G) are presented here.

(H) Lifetimes of FRET intermediates observed in indicated conditions in the FRET range from 0.35 to 0.5. See Supplemental Note 2 and Supplemental Table 2.

Supplemental Figure 2

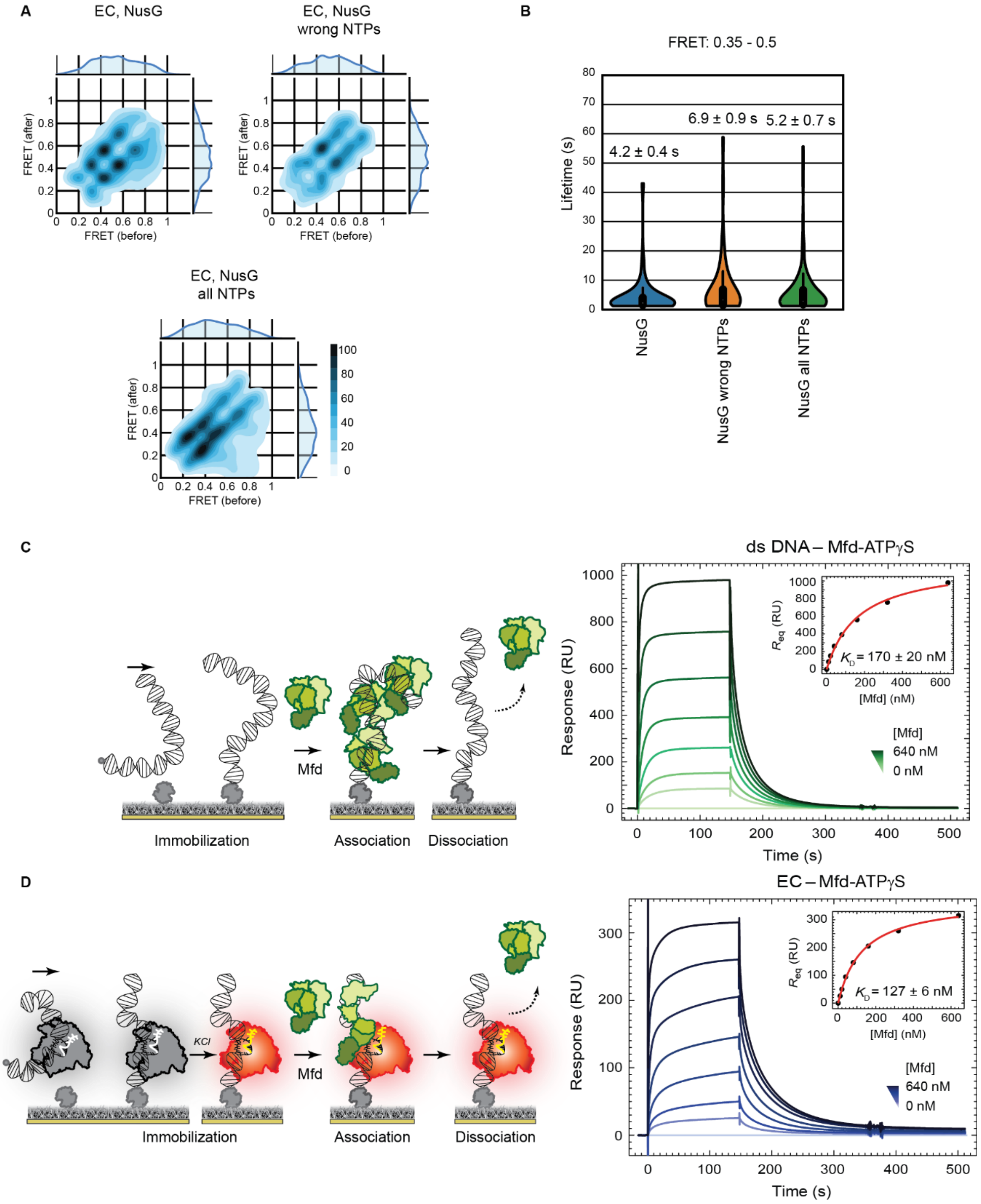

**Supplemental Figure 2: Measurement of binding affinity of Mfd for dsDNA and the EC.** ***Related to Figures 2 and 3.***

(A)Transition density plots for EC and NusG alone (n = 461 molecules), EC and NusG in the presence of wrong NTPs (n = 231 molecules), and EC and NusG with the full set of NTPs (n = 426 molecules).

(B) Lifetimes of FRET intermediates observed in indicated conditions in the FRET range from 0.35 to 0.5. See Supplemental Note 2 and Supplemental Table 2.

(C, D) Schematic and SPR sensorgrams show association (150 s) and dissociation phases of serially-diluted 10 – 640 µM Mfd-ATPɣS, including a 0 µM control for (C) biotinylated 49-mer dsDNA, and (D) *in vitro* reconstituted EC on biotinylated 49-mer DNA flowed into the SPR chip. Responses at equilibrium averaged over the steady region of sensorgrams ($R_{eq}$) were fit (insets) using a steady-state affinity (SSA) model (Equation 1, Materials and Methods) to derive values of $K_D$ (as indicated). Errors are S.E. of the fit.

### Supplemental Figure 3

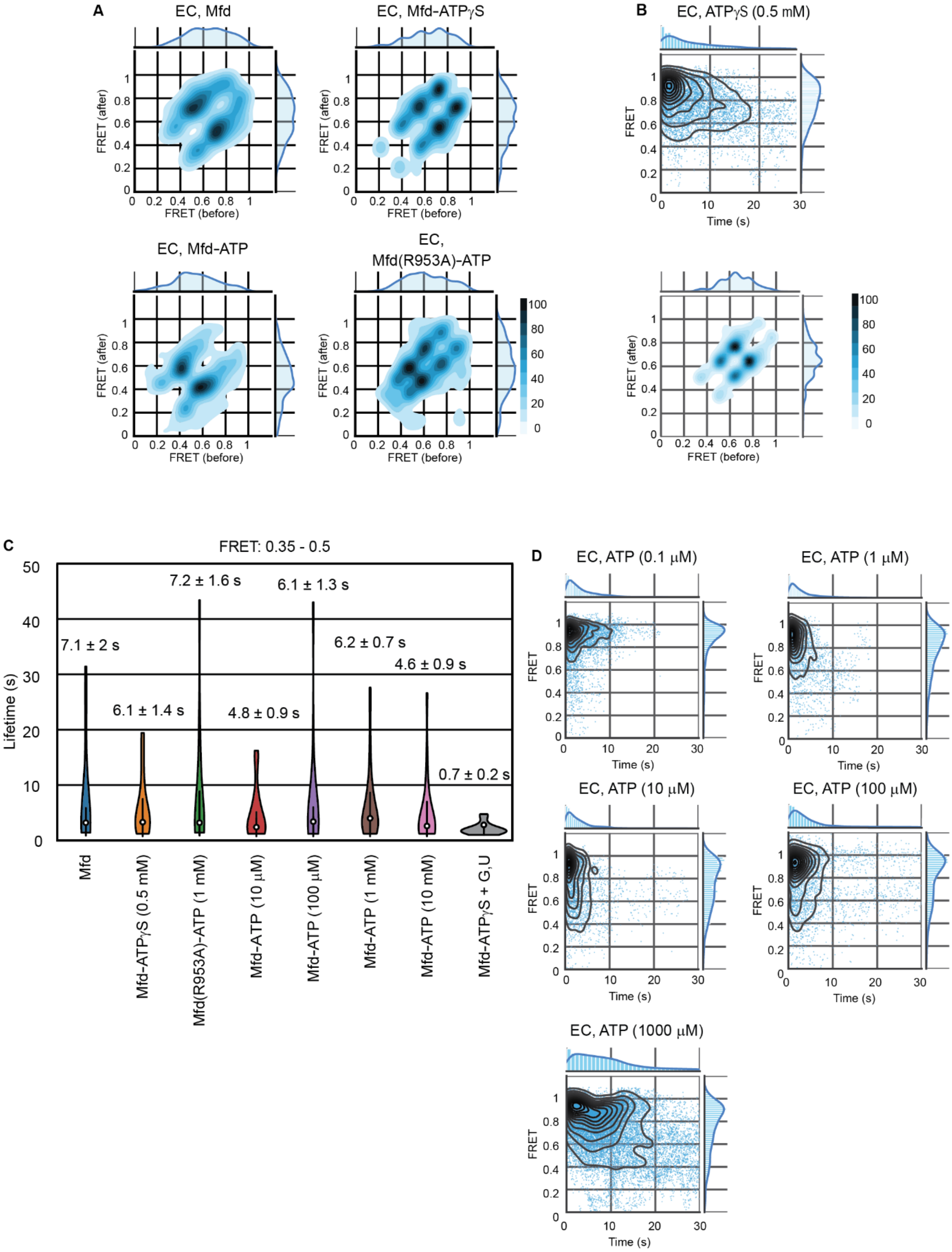

**Supplemental Figure 3: Interactions of ATP with the EC alone or in the presence of Mfd. *Related to Figure 3.***

(A) Transition density plots for EC and Mfd alone (n = 75 molecules), EC and Mfd in the presence of 1 mM ATPɣS (n = 68 molecules), EC and Mfd in the presence of 1mM ATP (n = 174 molecules), and EC and Mfd(R953A) in the presence of 1 mM ATP (n = 234 molecules).

(B) Temporal heat map (upper panel) and transition density plot (lower panel) of FRET pair labelled RNAP EC in the presence of 0.5 mM ATPɣS alone (n = 124 molecules).

(C) Lifetimes of FRET intermediates observed in indicated conditions in the FRET range from 0.35 to 0.5. See Supplemental Note 2 and Table 2.

(D) Temporal heat maps of FRET pair labelled RNAP EC when titrated with ATP in the range of 0.1 µM to 1 mM (n = 151, 82, 81, 76, 157 molecules respectively) as indicated.

### Supplemental Figure 4

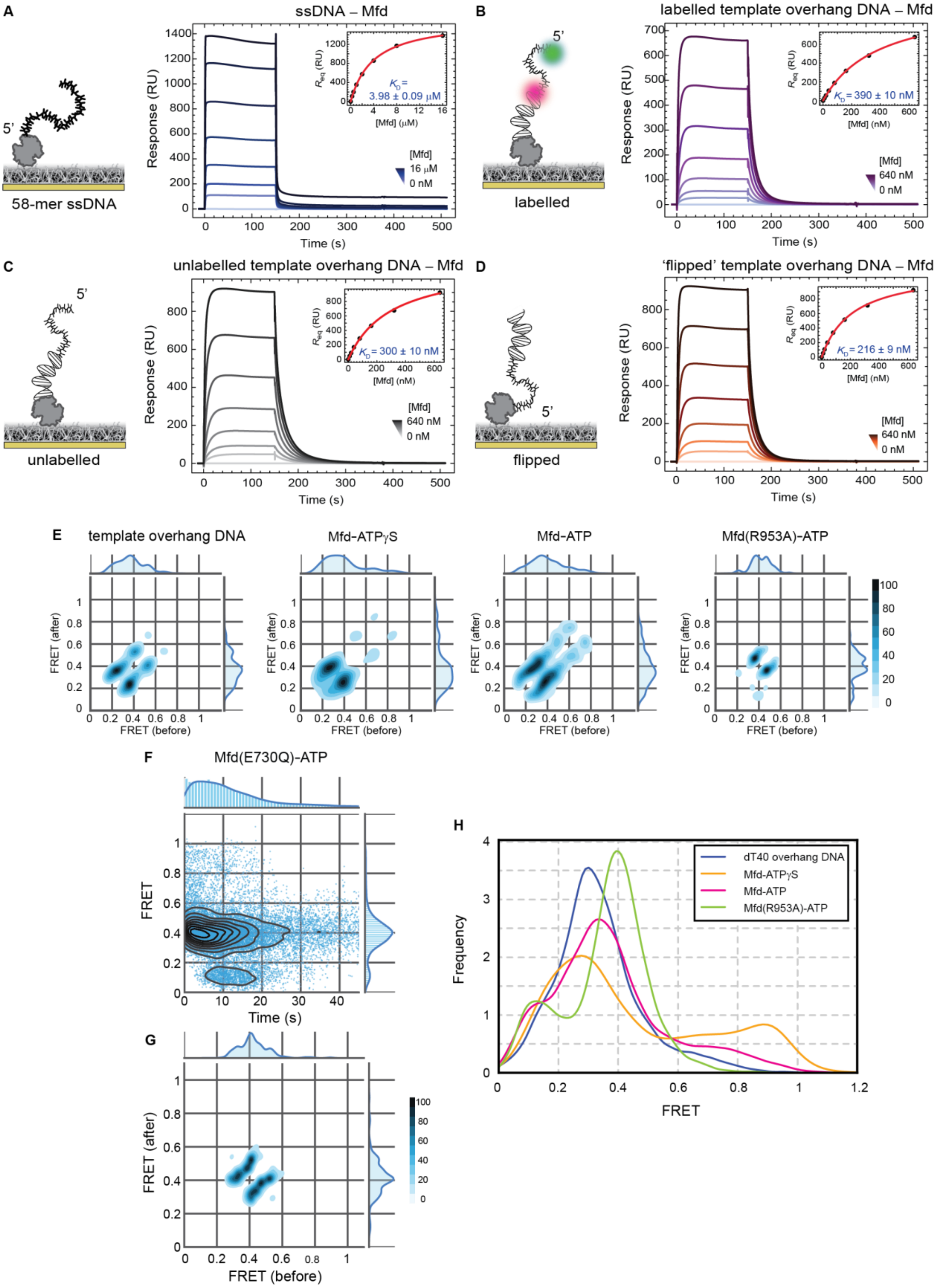

**Supplemental Figure 4: Interactions between Mfd and a primed-DNA template.** ***Related to Figure 4.***

Schematic and SPR sensorgrams show association (150 s) and dissociation phases of serially-diluted 10 – 640 µM Mfd-ATPɣS, including a 0 µM control for (A) 58-mer ssDNA containing the $dT_{40}$ sequence, (B) FRET pair labeled primed DNA substrate (biotinylated 18-mer dsDNA with $dT_{40}$ overhang), (C) primed DNA substrate (biotinylated 18-mer dsDNA with $dT_{40}$ overhang) and (D) “flipped” primed DNA substrate (18-mer dsDNA with biotinylated $dT_{40}$ overhang). Responses at equilibrium averaged over the steady region of sensorgrams ($R_{eq}$) were fit (insets) using a steady-state affinity (SSA) model to derive values of $K_D$ (as indicated). Errors are S.E. of the fit.

(E) Transition density plots for FRET-pair labeled primed DNA alone (n = 100 molecules), or bound to Mfd in the presence of ATPɣS (n = 94 molecules), or ATP (n = 203 molecules), and Mfd(R953A) in the presence of ATP (n = 105 molecules) are presented here. The DNA substrate is the same as described in Figure 4.

(F, G) Temporal heat map (F) and transition density plot (G) for ATPase mutant Mfd(E730Q) binding to FRET pair labelled primed DNA template used in Figure 4 (n = 103 molecules).

(H) For convenience, kernel density estimations of the ensemble FRET distributions read as projections along the ordinate in the heat-maps presented in Figure 4 are presented to enable direct visual comparisons.

Supplemental Figure 5

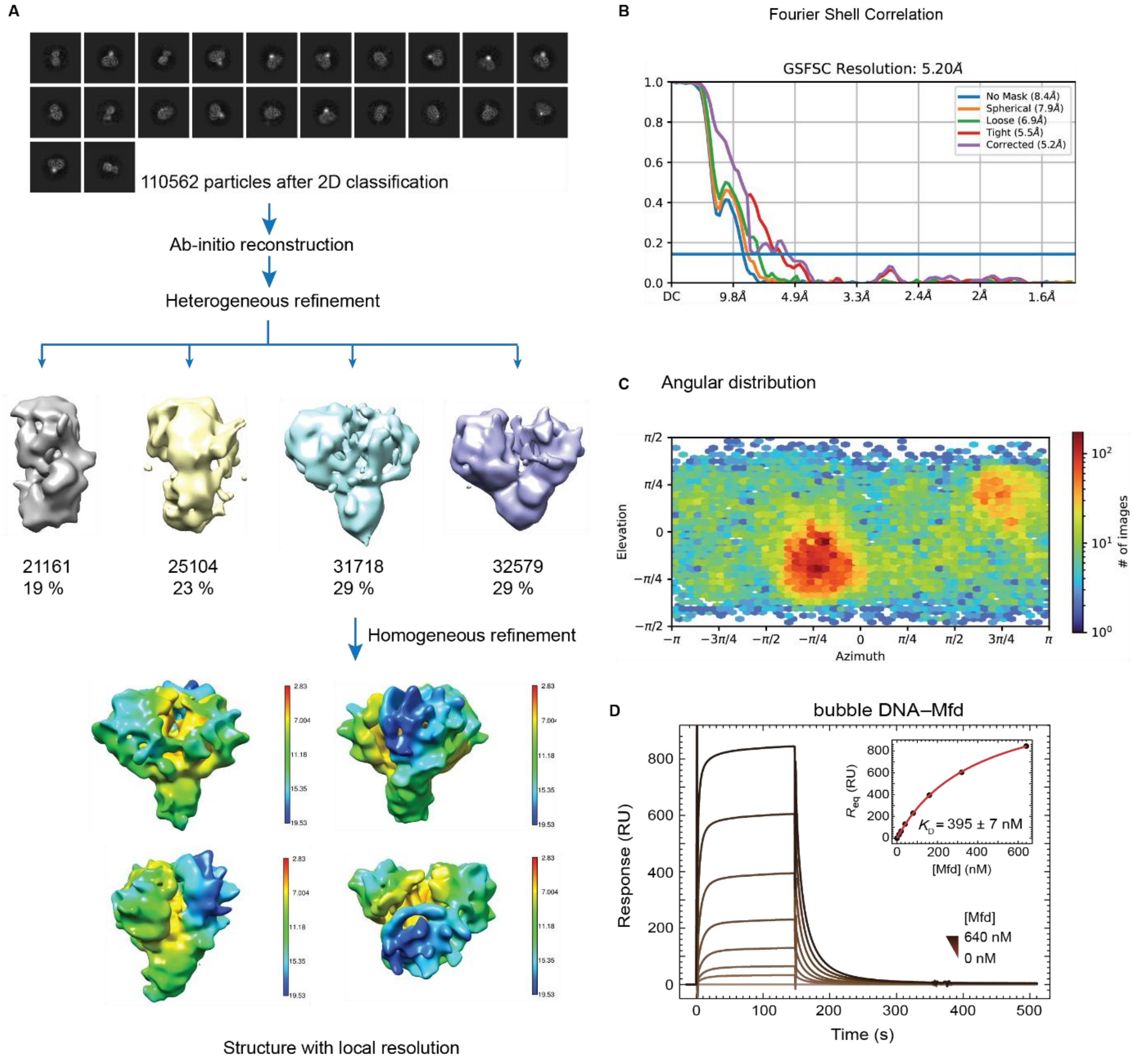

**Supplemental Figure 5: EM reconstruction of Mfd. *Related to Figure 5.***

(A) Overview of image processing using CryoSPARC. A set of 110562 particles were selected after 2D classification for *ab initio* reconstruction and heterogeneous refinement leading to four classes of 3D reconstructions. Of these, 31718 particles that yielded the most complete reconstruction were subjected to homogenous refinement to generate the map presented here. The colormap applied to the reconstruction represents local resolution.

(B) Fourier shell correlation curves of the reconstruction reported an apparent resolution of 5.2 Å (gold-standard 0.143 FSC).

(C) Angular distribution of particle projections

(D) Measurement of the dissociation constant of Mfd for a 49-mer dsDNA template containing a 9 nucleotide non-complementary bubble where the complementary residues were replaced by $dT_9$. The errors reflect S.E. of the fit.

**Supplemental Note 1**

To assess whether the calculated distances are consistent with the FRET state assignments for the pre- *vs*. post- translocated states, we built an *in silico* model (Supplemental Figure 1B) based on available structures of the *E. coli* elongation complex (Abdelkareem et al., 2019) and structures of the Cy3 and Cy5 dyes (Liu and Lilley, 2017). The distances between the Cy3 and Cy5 dyes in the pre- and post-translocated states were then estimated to be 1.7 nm and 3.9 nm respectively. Using these distances, and available estimates for $R_0$ for Cy3–Cy5 dye pairs introduced into protein-DNA complexes in the range of 4.7 – 6.5 nm (Levitus and Ranjit, 2011), we estimate FRET efficiencies for the pre- and post- translocated states to be: 0.99 and 0.75 (for $R_0$ = 4.7 nm) and 0.99 and 0.95 (for $R_0$ = 6.5 nm). The measured FRET efficiencies are ~0.9 and ~0.6 (Figure 1), which agree well with the lower estimates for $R_0$.

**Supplemental Note 2**

To measure the lifetime of the intermediate state we first chose to analyze trajectories with intermediates in the mid-FRET regime. However, examination of individual trajectories revealed that this mid-FRET regime is populated by molecules exhibiting high-mid FRET state dynamics as well as mid-low FRET dynamics. We therefore selected molecules in the 0.35–0.5 FRET range which mostly accommodated molecules exhibiting mid-low FRET dynamics. The mean lifetimes (and standard error) of the intermediates states were best estimated by fitting the distributions to single-exponential fits. In the case of one notable condition – Mfd in the presence of ATPɣS followed by incubation with correct nucleotides – the lifetime distributions was found to be better fit by a gamma distribution. In this case, the scale parameter was used to report the lifetime (Supplemental Table 2).

**Supplemental Note 3**

To compare the efficiency of transcription by the elongation complex following incubation with Mfd-ATPɣS upon addition of 100 µM each of GTP and UTP to that in the presence of GTP, UTP and ATP, we plotted histograms of the FRET states observed in the system (gray bars in Figure 3L). The histogram for each condition was best fit with a sum of three Gaussian terms of the form:

$$f(x) = ae^{-\frac{(x-\mu)^2}{2\sigma^2}}$$

Here $a_i$ represents the amplitude, $\mu_i$ represents the mean of the distribution and $\sigma_i$ the standard deviation of the $i^{\text{th}}$ distribution. In the absence of Mfd-ATPɣS, the three Gaussians $(a_i, \mu_i, \sigma_i)$ were: (25.36, 0.91, 0.11); (16.5, 0.63, 0.33) and (15.14, 0.09, 0.10) with an R-square of 0.88. In the presence of Mfd-ATPɣS and rGTP, rUTP, the three Gaussians were: (17.15, 0.93, 0.19), (10.94, 0.70, 0.27) and (9.82, 0.18, 0.10) with an R-square of 0.85. For each Gaussian the area was calculated as:

$$A_i = \sqrt{2\pi} a_i \sigma_i$$

Next, each area was normalized by the sum of the areas of the three Gaussians to obtain the fractional area occupied by each Gaussian. Efficiencies were calculated by comparing the fractions of the populations corresponding to each of the Gaussians across the two conditions. Since transcript elongation is reflected in the low-FRET states, we compared the fraction of the population in the Gaussian with the smallest mean FRET.

**Supplemental Note 4**

SPR measurements of the strength of interaction between Mfd and this DNA substrate in the presence of ATPɣS revealed a dissociation constant that is somewhat higher ($K_D$ = 300 ± 10 nM) than that of dsDNA (170 ± 20 nM; Supplemental Figure 4B–D compared to Supplemental Figure 2C). To monitor changes in the distance between the junction and the 5' end, we introduced a FRET pair on either end of the ssDNA overhang (Figure 4D). SPR measurements revealed an affinity of 390 ± 10 nM for the binding of Mfd to the FRET-pair labelled substrate (Supplemental Figure 4B).

**Supplemental Table 1: Nucleic Acid substrates used in this study**

| Experiment | DNA | Sequence |
|---|---|---|
| pETMCSII_Mfd_F | Cloning primer | GTT TAA TCG GAT CCT AAG GAG GTT AAT TCC CGC TAT GCC TGA ACA ATA TCG TTA TAC G |
| pETMCSII_Mfd_R | Cloning primer | GGGAGCTCGAATTCTTAAGCGATCGCGTTCTCT |
| smFRET (Figures 1–3) | Non-template (NT70_bio) | ATC GAG CAA CTA CTC AGA CAG CAC TAC TGC GAC TTA CAG ACA TCG AGA GGG TAA TGG CGA ATA GCA CTG A /3 bioTEG/ |
| smFRET (Figures 1–3) | Template (T70_31Cy3) | TCA GTG CTA TTC GCC ATT ACC CTC TCG ATG T/iCy3/CT GTA AGT CGC AGT AGT GCT GTC TGA GTA GTT GCT CGAT |
| smFRET (Figures 1–3) | RNA(R15_4Cy5) | rArUrArU/iCy5/rArU rArUrC rGrArG rArGrG |
| smFRET (Figure 4) | bio_AS18_Cy5 | /5Biosg/ TGG CGA CGG CAG CGA GGC/3Cy5Sp/ |
| smFRET (Figure 4) | Cy3_dT40_S18 | /5Cy3/TT TTT TTT TTT TTT TTT TTT TTT TTT TTT TTT TTT TTT TTG CCT CGC TGC CGT CGC CA |
| SPR (Supplemental Figure 4) | bio_AS18 | /5Biosg/ TGG CGA CGG CAG CGA GGC |
| SPR (Supplemental Figure 4) | dT40_S18 | TT TTT TTT TTT TTT TTT TTT TTT TTT TTT TTT TTT TTT TTG CCT CGC TGC CGT CGC CA |
| SPR (Supplemental Figure 4) | bio_dT40_S18 | /Biosg/TT TTT TTT TTT TTT TTT TTT TTT TTT TTT TTT TTT TTT TTG CCT CGC TGC CGT CGC CA |
| SPR | AS18 | TGG CGA CGG CAG CGA GGC |

| | | |
|---|---|---|
| (Supplemental Figure 4) | | |
| SPR (Supplemental Figure 2) | TS_49 | TCA GTG CTA TTC GCC ATT ACC CTC TCG ATG T/iCy3/CT GTA AGT CGC AGT AGT G |
| SPR (Supplemental Figure 2) | NT_49_bio | CAC TAC TGC GAC TTA CAG ACA TCG AGA GGG TAA TGG CGA ATA GCA CTG A /3 bioTEG/ |
| SPR (Supplemental Figure 5) | NT_49_bio_ bubble | CAC TAC TGC GAC TTA CAG ACT TTT TTT TTG TAA TGG CGA ATA GCA CTG A /3BioTEG/ |
| SPR (Supplemental Figure 2) | RNA_15 | rArUrAr UrArU rArUrC rGrArG rArGrG |
| SPR (Figure 4) | S18 | GCC TCG CTG CCG TCG CCA |
| SPR (Figure 4) | dT3_S18 | TTT GCC TCG CTG CCG TCG CCA |
| SPR (Figure 4) | dT6_S18 | TTT TTT GCC TCG CTG CCG TCG CCA |
| SPR (Figure 4) | dT9_S18 | TTT TTT TTT GCC TCG CTG CCG TCG CCA |
| SPR (Figure 4) | dT12_S18 | TTT TTT TTT TTT GCC TCG CTG CCG TCG CCA |
| SPR (Figure 4) | dT15_S18 | TTT TTT TTT TTT TTT GCC TCG CTG CCG TCG CCA |
| SPR (Figure 4) | S18_bio | GCC TCG CTG CCG TCG CCA /3Bio/ |
| SPR (Figure 4) | AS18_dT3 | TGG CGA CGG CAG CGA GGC TTT |

| | | |
|---|---|---|
| SPR (Figure 4) | AS18_dT6 | TGG CGA CGG CAG CGA GGC TTT TTT |
| SPR (Figure 4) | AS18_dT9 | TGG CGA CGG CAG CGA GGC TTT TTT TTT |
| SPR (Figure 4) | AS18_dT12 | TGG CGA CGG CAG CGA GGC TTT TTT TTT TTT |
| SPR (Figure 4) | AS18_dT15 | TGG CGA CGG CAG CGA GGC TTT TTT TTT TTT TTT |
| 2AP Bulk fluorescence (Figure 5) | dT6_S18_v2_11_2AP | TTT TTT GAC T/i2AmPr/A CAG CCG ACG CGT |
| 2AP Bulk fluorescence (Figure 5) | dT6_S18_v2_12_2AP | TTT TTT GAC TA/i2AmPr/ CAG CCG ACG CGT |
| 2AP Bulk fluorescence (Figure 5) | AS18_v2_2AP | ACG CGT CGG CTG TTA GTC |
| 2AP Bulk fluorescence (Figure 5) | AS18_v2_2AP_bubble | ACG CGT CGG CTG AAA GTC |

**Supplemental Table 2: Lifetimes of mid-FRET states**

| | Condition | Distribution | Fit Parameter | | | | |
|---|---|---|---|---|---|---|---|
| | | | Mean (τ) [Exp]/ scale parameter (Gamma) | S.E. | Shape | S.E. | log likelihood |
| 1 | EC (n_states = 42) | exp | 5.0 | 0.8 | | | –109.9 |
| 2 | EC dNTPs (n_states = 14) | exp | 5.7 | 1.5 | | | –38.4 |
| 3 | EC wrong (n_states = 70 | exp | 7.8 | 0.9 | | | –213.7 |
| 4 | EC right (n_states = 19) | exp | 5.5 | 1.3 | | | –51.6 |
| 5 | NusG (n_states = 99) | exp | 4.2 | 0.4 | | | –242.1 |
| 6a | NusG wrong (n_states = 57) | exp | 6.9 | 0.9 | | | –166.9 |
| 7a | NusG correct (n_states = 129) | exp | 5.9 | 0.5 | | | –356.965 |
| 8 | Mfd (n_states = 13) | exp | 7.1 | 2.0 | | | –38.5 |
| 9 | Mfd-ATPɣS (n_states = 20) | exp | 6.1 | 1.4 | | | –56.1 |
| 10 | Mfd(R953A) (n_states = 21) | exp | 7.2 | 1.6 | | | –62.3 |
| 11 | Mfd-ATP [10µM] (n_states = 29) | exp | 4.8 | 0.9 | | | –74.9 |
| 12 | Mfd-ATP [100µM] (n_states = 23) | exp | 6.1 | 1.3 | | | –64.5 |

| | | | | | | | |
|---|---|---|---|---|---|---|---|
| 13 | Mfd-ATP [1 mM]<br><br>(n_states = 77) | exp | 6.2 | 0.7 | | | –216.9 |
| 14 | Mfd-ATP [10 mM]<br><br>(n_states = 27) | exp | 4.6 | 0.9 | | | –68.1 |
| 15 | ATPɣS_rG_rU<br><br>(n_states = 17) | gamma | 0.7 | 0.2 | 5.20 | 1.73 | –29.6 |
